# DE Novo emerged gene SEarch in Eukaryotes with DENSE

**DOI:** 10.1101/2024.01.30.578014

**Authors:** Paul Roginski, Anna Grandchamp, Chloé Quignot, Anne Lopes

## Abstract

The discovery of de novo emerged genes, originating from previously noncoding DNA regions, challenges traditional views of species evolution. Indeed, the hypothesis of neutrally evolving sequences giving rise to functional proteins is highly unlikely. This conundrum has sparked numerous studies to quantify and characterize these genes, aiming to understand their functional roles and contributions to genome evolution. Yet, no fully automated pipeline for their identification is available. Therefore, we introduce DENSE (DE Novo emerged gene Search), an automated Nextflow pipeline based on two distinct steps: detection of Taxonomically Restricted Genes (TRGs) through phylostratigraphy, and filtering of TRGs for de novo emerged genes via genome comparisons and synteny search. DENSE is available as a user-friendly command-line tool, while the second step is accessible through a web server upon providing a list of TRGs. Highly flexible, DENSE provides various strategy and parameter combinations, enabling users to adapt to specific configurations or define their own strategy through a rational framework, facilitating protocol communication, and study interoperability. We apply DENSE to seven model organisms, exploring the impact of its strategies and parameters on de novo gene predictions. This thorough analysis across species with different evolutionary rates reveals useful metrics for users to define input datasets, identify favorable/unfavorable conditions for de novo gene detection, and control potential biases in genome annotations. Additionally, predictions made for the seven model organisms are compiled into a requestable database, that we hope will serve as a reference for de novo emerged gene lists generated with specific criteria combinations.

**Significance Statement:** The identification and classification of de novo genes, which originate from noncoding regions of DNA, remain an ongoing challenge in genomics research. While various approaches have been employed for their identification, the lack of a standardized protocol has resulted in varying lists of de novo genes across studies. This study introduces a novel tool: DENSE, that formalizes the common practices used in the field into a comprehensive and automated pipeline. DENSE streamlines the identification of taxonomically restricted genes, homology searches, and synteny analysis. This standardized methodology aims to enhance the accuracy and reliability of de novo gene identification, fostering a deeper understanding of the evolutionary mechanisms that drive gene birth and shape the genetic diversity of organisms.

## Introduction

Comparative genomics has unveiled the existence of what we call de novo emerged genes – genes that arose from a DNA region that was ancestrally noncoding. Initially considered unlikely (Jacob, 1977), the accumulation of sequencing data revealed that they were in fact, widespread, being detected in various eukaryotic species, and numerous, with several dozens of examples reported for different organisms (Cai et al., 2008; Levine et al., 2006; Schlötterer, 2015; Van Oss & Carvunis, 2019). The discovery of these genes has highlighted that the passage from the noncoding world to the world of functional products and regulated processes was much more frequent than previously thought, challenging previous assumptions and raising an intriguing question: How can a noncoding neutrally-evolving DNA sequence give rise to a functional product able to take part in the well-established biological networks and contribute to the organism’s fitness? Indeed, noncoding regions are associated with different nucleotide compositions and combinations from coding regions and are consequently expected to result in nonfunctional combinations of amino acids if translated. The newly attributed role to noncoding regions and the unexpected permeability observed between these two worlds have thus captured the attention of researchers in recent years. Notably, numerous studies have been undertaken on different species, including several model organisms, to characterize de novo emerged genes (Bungard et al., 2017; Lange et al., 2021; Papadopoulos et al., 2021; Peng & Zhao, 2023; Schmitz et al., 2018; Vakirlis et al., 2018). This effort not only aims to elucidate their properties and functions but also to serve as a proxy for a better understanding of the transition between the noncoding and the coding worlds as well as their evolutionary relationship. The strategy employed to detect de novo emerged genes being likely to influence the resulting interpretations, it is therefore important to design rational and reproducible methods.

So far, most model organisms have been associated with multiple lists of de novo emerged genes as illustrated in the Table 1 of the review of Oss and Carvunis (Van Oss & Carvunis, 2019). These lists result from diverse protocols that generally rely on transcriptomics and/or comparative genomics. Nonetheless, despite this diversity, all these methods, for classifying genes as de novo emerged, impose that genes be novel or identified in closely related species. Transcriptomics-based protocols either search for novel transcripts or require the transcription of de novo gene candidates usually detected with an initial step of comparative genomics (Blevins et al., 2021; Grandchamp et al., 2023; Zhang et al., 2019). However, all novel transcripts, especially those resulting from pervasive expression, are not necessarily expected to last throughout evolution and can instead be considered as gene precursors (Grandchamp et al., 2023; Wacholder et al., 2023). As discussed in previous reports, transcription may predate gene emergence and, therefore, cannot be considered a sufficient criterion for a novel transcript to be classified as a gene (Cai et al., 2008; Carvunis et al., 2012; Chen et al., 2020; Papadopoulos et al., 2023; Reinhardt et al., 2013; Schlötterer, 2015). Furthermore, young de novo emerged genes are typically associated with stress response or adaptation, and are expected to be expressed under specific conditions (Carvunis et al., 2012; Colbourne et al., 2011; Donoghue et al., 2011; Schlötterer, 2015; Van Oss & Carvunis, 2019). Thus, demonstrating the expression of such genes involves finding the conditions under which they are expressed, which is not trivial. Consequently, requiring de novo gene candidates to be transcribed may be accompanied by high ratios of false negatives.

**Table 1:**
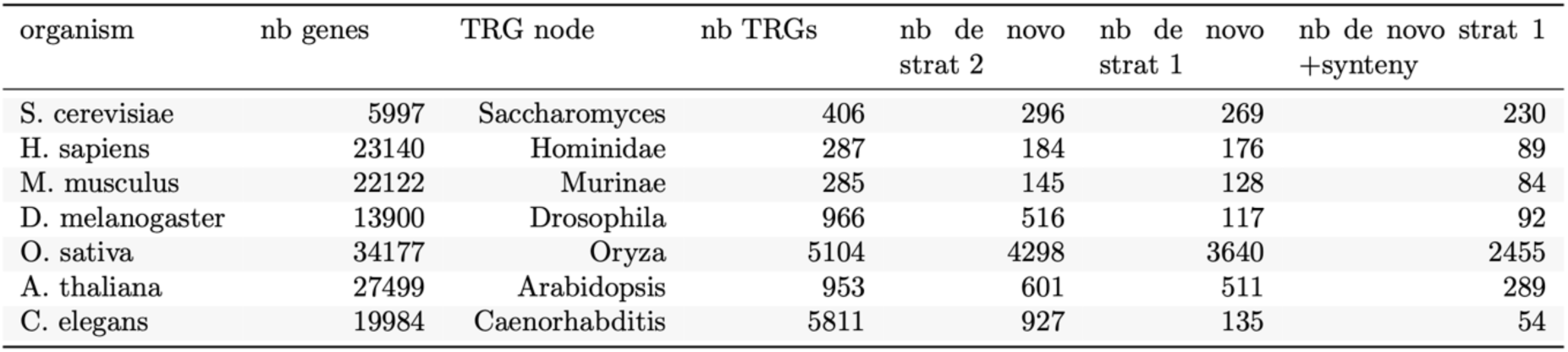
DENSE predictions for the seven studied species: For each species, are indicated its number of genes, the phylostratum threshold used for TRG detection, the number of predicted TRGs, and the number of de novo emerged genes predicted with Strategy 2, Strategy 1 or Strategy combined with synteny criterion.

| organism | nb genes | TRG node | nb TRGs | nb de novo<br>strat 2 | nb de novo<br>strat 1 | nb de novo strat 1<br>+synteny |
| --- | --- | --- | --- | --- | --- | --- |
| *S. cerevisiae* | 5997 | Saccharomyces | 406 | 296 | 269 | 230 |
| *H. sapiens* | 23140 | Hominidae | 287 | 184 | 176 | 89 |
| *M. musculus* | 22122 | Murinae | 285 | 145 | 128 | 84 |
| *D. melanogaster* | 13900 | Drosophila | 966 | 516 | 117 | 92 |
| *O. sativa* | 34177 | Oryza | 5104 | 4298 | 3640 | 2455 |
| *A. thaliana* | 27499 | Arabidopsis | 953 | 601 | 511 | 289 |
| *C. elegans* | 19984 | Caenorhabditis | 5811 | 927 | 135 | 54 |

On the other hand, comparative genomics appears as an appealing solution since they can be applied once the complete genome of the species of interest (i.e., the focal species) and those of several of its close neighboring species are available. Although these methods may involve more or less stringent criteria, all of them focus on genes that have emerged recently, and first rely on the detection of taxonomically-restricted genes (TRGs) (Peng & Zhao, 2023; Vakirlis et al., 2018; Vakirlis & McLysaght, 2019; Weisman, 2022). The latter correspond to genes found in a single species (i.e., orphan genes) or closely related species, a prerequisite for finding young de novo emerged genes. The detection of TRGs generally relies on phylostratigraphy that estimates the age of a gene of interest from the detection of its homologs across a phylogeny (Arendsee, Li, Singh, Seetharam, et al., 2019; Barrera-Redondo et al., 2023; Domazet-Loso et al., 2007). In practice, the idea is to screen each gene of the focal genome against a large sequence database (typically, nr or uniprot) with BLAST or an equivalent tool (Altschul et al., 1990; Buchfink et al., 2021). The genes are then assigned an evolutionary age corresponding to the common ancestor of all the lineages in which they have been detected, i.e., the most distant node in the phylogeny where BLAST has detected a homolog of the considered gene. As such, it becomes evident that the criteria used for the homology search are critical and deserve to be defined with caution (Domazet-Lošo et al., 2017; Moyers & Zhang, 2015). Indeed, a lack of sensitivity would be accompanied by an underestimation of gene age, thereby wrongly annotating old genes as TRGs. However, thorough analyses that aimed to assess the impact of BLAST criteria on gene age estimation have demonstrated that the trends deduced from phylostratigraphy-based approaches remain robust in the face of BLAST’s lack of sensitivity, with some studies reporting optimal E-value thresholds in the context of TRG detection (Domazet-Loso et al., 2007; Domazet-Lošo et al., 2017; Vakirlis et al., 2020).

Finally, TRGs consist of a heterogeneous population and encompass genes with different origins, including genes resulting from duplication or horizontal transfer followed by high divergence. Ideally, one would aim to reconstruct the ancestral noncoding sequence to retrace the evolutionary stages that led to de novo gene birth. Since it is not always possible, (it requires the detection of an outgroup noncoding sequence with respect to the ancestor to be predicted), different approaches have been described in the literature to discriminate de novo emerged genes from the other TRGs (Peng & Zhao, 2023; Vakirlis & McLysaght, 2019; Van Oss & Carvunis, 2019; Weisman, 2022). Specifically, inferring that the locus of the gene candidate was noncoding in the ancestor through the analysis of the sister lineages offers an attractive alternative, providing solid support for de novo gene birth. Methods therefore require the identification of homology traces of the gene of interest in the noncoding regions of a species where the gene is absent. However, some of them require additional constraints to further support the noncoding status of the corresponding locus in the ancestor. Typically, the homology traces may be imposed to be found in a noncoding region of what we refer to as outgroup species ((Peng & Zhao, 2023; Weisman, 2022; Zhang et al., 2019). The latter are defined as those for which the gene is absent and that branch in the tree after the last species where the gene is present. In addition, to further assert the ancestor’s noncoding status, synteny-based approaches may be employed to guarantee the correct identification of the orthologous noncoding region. The latter therefore search for the syntenic region in the outgroup species and verifies the noncoding status of the homologous region (Knowles & McLysaght, 2009; McLysaght & Hurst, 2016; Tautz & Domazet-Lošo, 2011; Van Oss & Carvunis, 2019). Doing so, one may anticipate removing random homology signals expected to be more frequent with TRGs that are generally shorter than canonical coding sequences (CDSs). Various strategies have been undertaken for the identification of syntenic regions (Arendsee, Li, Singh, Bhandary, et al., 2019; Armstrong et al., 2020; Elghraoui et al., 2023)). If synteny blocks can be readily detected within very closely related species with high-quality genome assemblies, this task becomes difficult as the evolutionary distance between species increases, when studying genomes associated with high rates of chromosomal rearrangements or simply, when dealing with multiple contig genome assemblies (Liu et al., 2018; Ranz et al., 2001). Microsynteny, which searches for local gene order, nevertheless, offers a good compromise (Vakirlis & McLysaght, 2019). Beyond the fact that we have no *a priori* on the recombination rates in the regions that constitute hotbeds for de novo gene birth, the use of microsynteny enables one to handle genome assemblies of intermediate quality, therefore extending the applicability of such methods to wider genomic contexts.

Although a methodological consensus in comparative genomics-based approaches seems to have emerged in recent years, to the best of our knowledge, the scientific community still lacks a fully automated pipeline, different combinations of criteria and parameters are still reported, and no definitive protocol has yet been established. This hinders reproducibility and fair comparisons between studies, yet essential to decipher and eventually reconcile contradictory trends. Therefore, we propose DENSE (De Novo emerged gene SEarch), an automated pipeline that handles the entire process of de novo emerged gene detection, from identifying TRGs to filtering for those likely to have emerged de novo (https://github.com/Proginski/dense). Since protocols may continue to evolve, but also due to the heterogeneity in the quality of genome assemblies and/or annotations among species, DENSE has been designed to be highly flexible and offers various combinations of filters and parameters embedded in a unified Nextflow framework (Di Tommaso et al., 2017). In this manuscript, we introduce DENSE and investigate the impact of its different implemented strategies, as well as the influence of input data, on the prediction of de novo emerged genes. Finally, we present several metrics that can help users define their input dataset, identify favorable/unfavorable conditions for the detection of de novo emerged genes, and control for potential bias in genome annotations.

## Results

### Principle of DENSE

DENSE consists of two main independent steps that respectively search for TRGs among the annotated genes of a focal genome and then, identifies through a cascade of filters, those that have homology traces in the orthologous region of a species where the gene is absent with more or less stringent criteria (Figure 1). Specifically, DENSE starts from the genome of the focal species and those of its neighboring species along with their corresponding phylogenetic tree. Then based on the phylostratigraphy calculated by GenEra (Barrera-Redondo et al., 2023), it predicts the date of emergence of all annotated coding sequences (CDSs) of the focal genome with the assumption that horizontal transfers are rare in eukaryotic species. To do so, GenEra screens each CDS against the nr database and the annotated CDSs of the neighboring genomes with DIAMONDv2 (Buchfink et al., 2021)(Figure 1A). Alternatively, users have the option to screen other databases, such as uniprot, swissprot or a custom database. Nevertheless, despite being time-consuming for large genomes, we recommend using nr for more accurate age estimation. Each CDS is then assigned an evolutionary age, corresponding to the most distant node in the NCBI phylogeny where DIAMONDv2 has detected the CDS. In cases where a gene has multiple isoforms, the age of the oldest one is assigned to all isoforms. DENSE provides the user with all CDSs associated with their predicted evolutionary age and the list of the TRGs of the focal species (Figure 1B). One should note that, by default, the phylostratum level used to define TRGs is set to the genus level, but, depending on the studied species and the aims of the user, DENSE offers the possibility to modify it to younger or older phylostrata. To limit the number of false positives in TRG detection, DIAMONDv2 is called with the sensitive mode with a threshold E-value of 10^−5^ that has been shown to be optimal for the identification of orphan genes in *S. cerevisiae*, *D. melanogaster* and *H. sapiens* (Vakirlis et al., 2020).

**Fig. 1:**
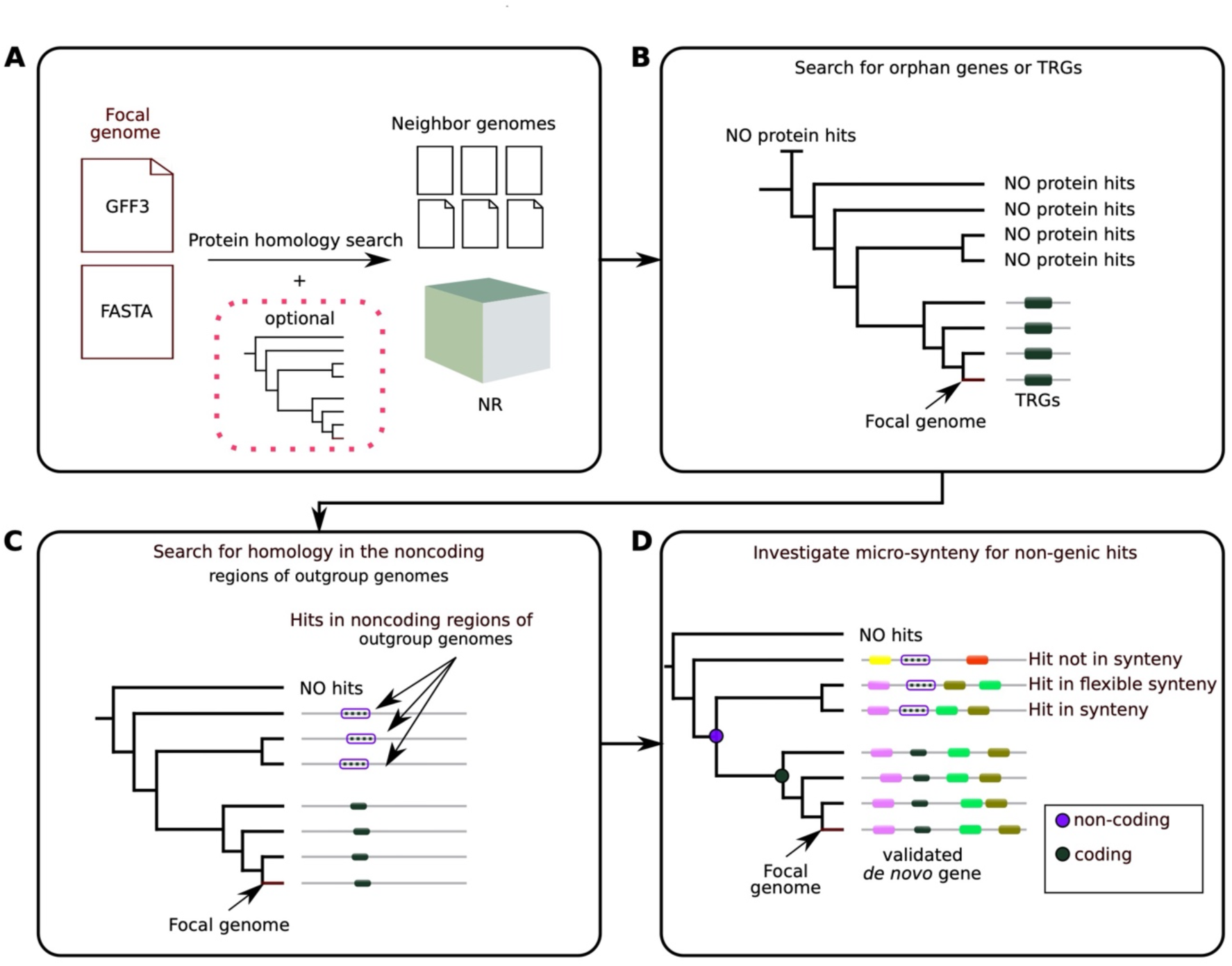
DENSE workflow. **a**) DENSE, through GenEra (Barrera-Redondo et al., 2023) screens all the focal genome’s annotated CDSs against the nr database and the genomes of the focal’s neighbors. Depending on the chosen strategy (mandatory for strategies 1 and 3), the tree of the focal genome and its neighbors (local tree) must be provided. Then DENSE outputs the estimated ages of each CDS of the focal genome and extracts its TRGs, including orphans, according to the phylostratum threshold indicated by the user (genus level by default). **b)** Example of the conservation of a focal TRG across the local tree. If no local tree has been provided, only the presence/absence across the neighbors is considered. **c)** Here, we assume that Strategy 1 has been called. DENSE screens the noncoding regions of the outgroup species (only applicable for the focal’s neighbors) with tblastn. In this example, three genomes are associated with a tblastn hit in a noncoding region (dashed lines surrounded by purple borders). **d)** We assume that the user requires the nonocoding hits to be found in synteny with the de novo emerged gene candidate. Therefore, DENSE verifies that at least one of the noncoding hits of the outgroup species is effectively detected in a region that is syntenic to that of the candidate. If so, the corresponding locus of their associated MRCA is considered as noncoding while that of the MRCA of the lineages where the candidate has been detected is considered as coding.

During the second step, DENSE focuses on the focal genome and the neighboring genomes provided by the user to apply a combination of filters in order to distinguish de novo emerged genes from the TRGs identified by GenEra in the initial step. Users can choose from several strategies, each associated with specific filter and parameter combinations (Figure 2A). Here, we present one of the strictest strategies (i.e., Strategy 1 combined with a search of synteny) that involves using tblastn for each TRG to search for sequence homology in the noncoding regions of the outgroup species (Figure 1CD, Figure 2B)(Gertz et al., 2006). Then, using microsynteny, DENSE checks whether the noncoding hit(s) detected in the outgroup species are located in synteny with the focal de novo gene candidate (Figure 1D, Figure 2C). Therefore, DENSE defines two gene windows of a specific size (set to four genes by default), flanking the focal gene (focal windows) and its homologous noncoding hit (target windows). It then requires at least one upstream gene and one downstream gene from the focal windows (number of anchor pairs set to one) to be present in the target windows. To do so, DENSE employs the Best Reciprocal Hit method (E-value = 10^−3^, and query coverage = 70%) to search for the orthologs of the focal windows within the outgroup neighbors. It then verifies the presence of at least two of these orthologs in the target windows, with one in the upstream window and the other in the downstream one (Figure 2C). One should notice that the number of anchor gene pairs and the size of the windows are parameterizable and that synteny search can be combined to any of the three strategies available in DENSE. Finally, all the TRGs that fulfill these criteria, (i.e., one hit in a syntenic noncoding region of at least one outgroup species), are considered genes that have recently emerged de novo (i.e., de novo gene candidates). Nevertheless, DENSE is very flexible and users can provide their own list of TRGs and enter the pipeline directly at the filtering step (Figure 1C) or define their own criteria. Notably, users can deactivate the verification of microsynteny and/or focus on candidates with hits in the noncoding regions of the neighboring species regardless of the fact that the species are outgroup. If removing filters may be expected to be accompanied by increases in false positive rates, this can be nevertheless useful when the phylogeny of the considered species is unknown, or incomplete, or when the outgroup species are too far to detect homology traces in their noncoding regions.

**Fig. 2:**
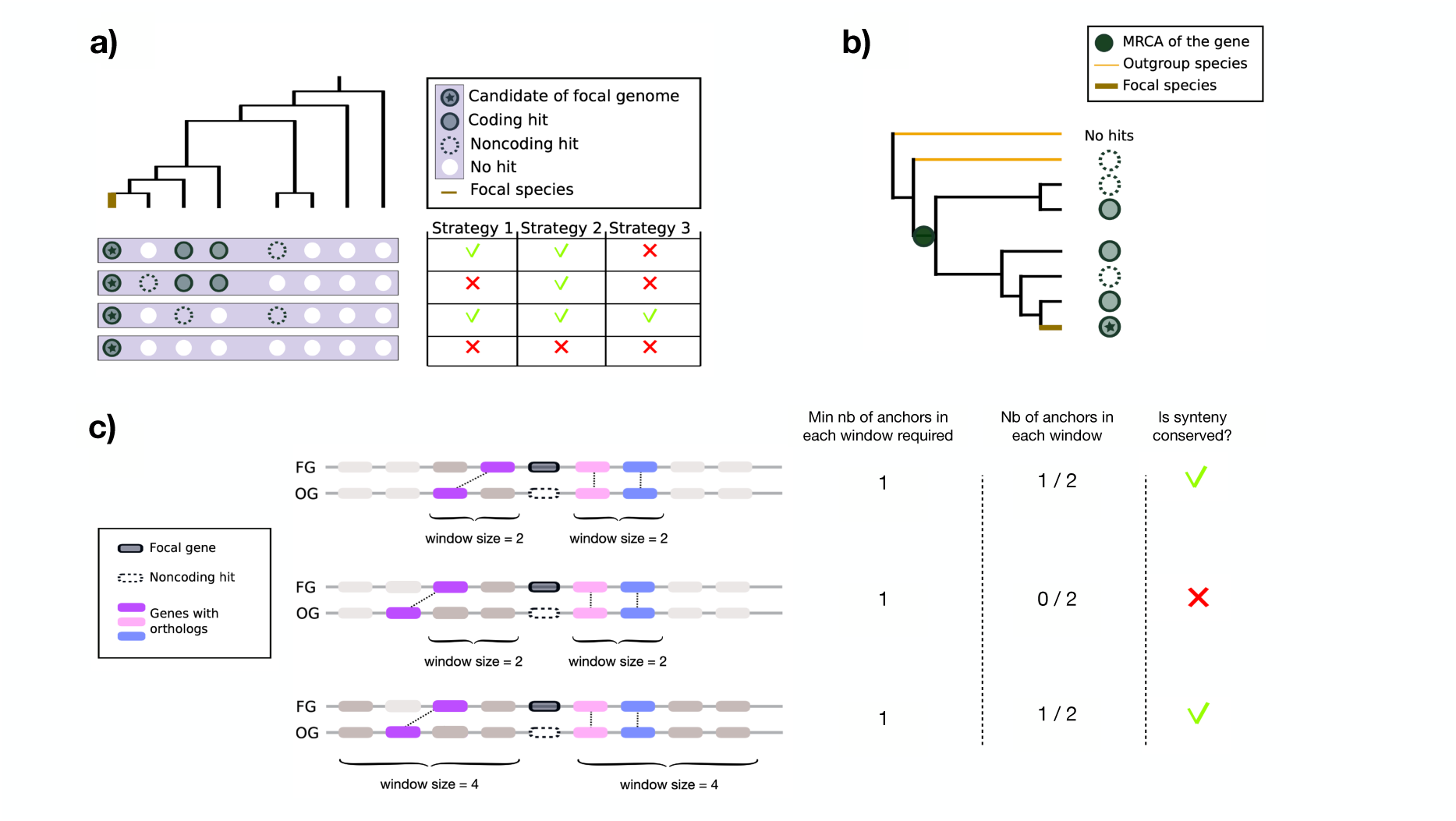
DENSE strategies and definitions. **a**) Strategies proposed by DENSE. Strategy 1 requires a noncoding hit in at least one outgroup species, while Strategy 2 only asks the candidate to have a noncoding hit, irrespective of the tree location of the corresponding species. Strategy 3 is the most stringent one since it requires candidates retained by Strategy 1 to be orphans with a noncoding hit. Here, are represented the absence/presence of coding and noncoding hits of four genes and their predicted status according to the employed strategy. Note that all strategies can be combined to the additional requirement of synteny conservation. **b)** Definition of what is referred to as an outgroup species in DENSE for a given gene. Same tree as panel a), with the focal species represented in brown. The absence/presence of coding and noncoding hits of the gene is represented on the right of the tree following the same scheme as in panel a). The orange node corresponds to the MRCA of the lineages where the gene is present and all species that branch after this node are considered as outgroup species according to this gene. **c)** Examples of synteny conservation checking for a de novo gene candidate (in black) of a focal genome (FG) and its homologous noncoding hit (in white, dashed borders) in an outgroup genome (OG). DENSE defines two windows of the same size located upstream and downstream of the focal gene (focal window) and its noncoding counterpart (target window). In all examples, at least one gene from each focal window must be detected in the target windows, with one gene located in the upstream and the other in the downstream target window (i.e., number of anchor pairs set to one). In the first example, one of the two genes of the upstream focal window is retrieved in the upstream target window (purple orthologous pair forming the upstream anchor), while the two genes of the downstream focal window are retrieved in the downstream target window (orthologous pairs in pink and blue forming the downstream anchors). The synteny is then considered as conserved since each window is associated with at least one anchor gene. In the second example, no anchor is found within the upstream window (i.e., the purple ortholog is located outside the target window), thus the conservation of synteny is not validated. In the last example, the window size is extended to four genes, enabling the purple ortholog in the outgroup species to be retrieved. The two focal windows are associated with an anchor, thus the conservation of synteny is validated.

### Application of DENSE to seven model organisms

To illustrate the usage of DENSE, we applied it to seven model organisms that are well annotated and for which at least four neighboring species were sequenced and annotated (see list in Table 1, and complete list of neighbors in Supplementary Table S1). We used the default parameters, which require genes to have a tblastn hit in a syntenic noncoding region of an outgroup species, to be predicted as de novo emerged (Strategy 1 with synteny, using a window of four genes with one anchor pair). With the exception of *O. sativa*, we detected several dozen to a few hundred de novo gene candidates in all model organisms (Table 1 for the number of candidates detected at each step). The majority of detected de novo gene candidates, irrespective of the focal species, are orphans (68%)(Figure 3) or identified in the closest neighbors, illustrating the difficulty of detecting events of de novo gene emergence older than several million years ago. Interestingly, DENSE predicted approximately 2500 de novo emerged genes for *O. sativa*. Most of them (77%) are also very young, exclusively detected within *O. sativa*, despite the close proximity of its nearest neighbors which are less than 1.5 million years distant (Figure 3). In fact, this result must be interpreted with caution as it does not necessarily imply a higher propensity for de novo gene birth in *O. sativa* compared to the other species, and may result from methodological reasons. To classify a gene as de novo emerged, DENSE needs to detect for each de novo gene candidate, a tblastn hit in the syntenic noncoding region of an outgroup species (Figure 1D). This becomes more challenging as the distance between the focal and the outgroup species increases. Indeed, noncoding regions evolve fast, and the detection of homology relationships with such fast-evolving sequences is expected to decrease rapidly with the evolutionary distance separating the focal and the screened neighbors. Consequently, our ability to confidently support a TRG as a de novo emerged gene is bounded by our capacity to detect its orthologous noncoding hit(s) in the outgroup species, which in turn, is directly limited by the distance of the latter. Precisely, *O. sativa* has seven neighbors with less than 1.5 million years of divergence, which probably facilitates the detection of outgroup noncoding hits. In contrast, the closest neighbors of the other focal species (except *A. thaliana*) are more than 4 million years distant according to TimeTree (Kumar et al., 2022). However, it is noteworthy that geological time is not well-suited for fair comparisons between such diverse species. Each species is characterized by different generation times and may evolve at distinct evolutionary rates.

**Fig. 3:**
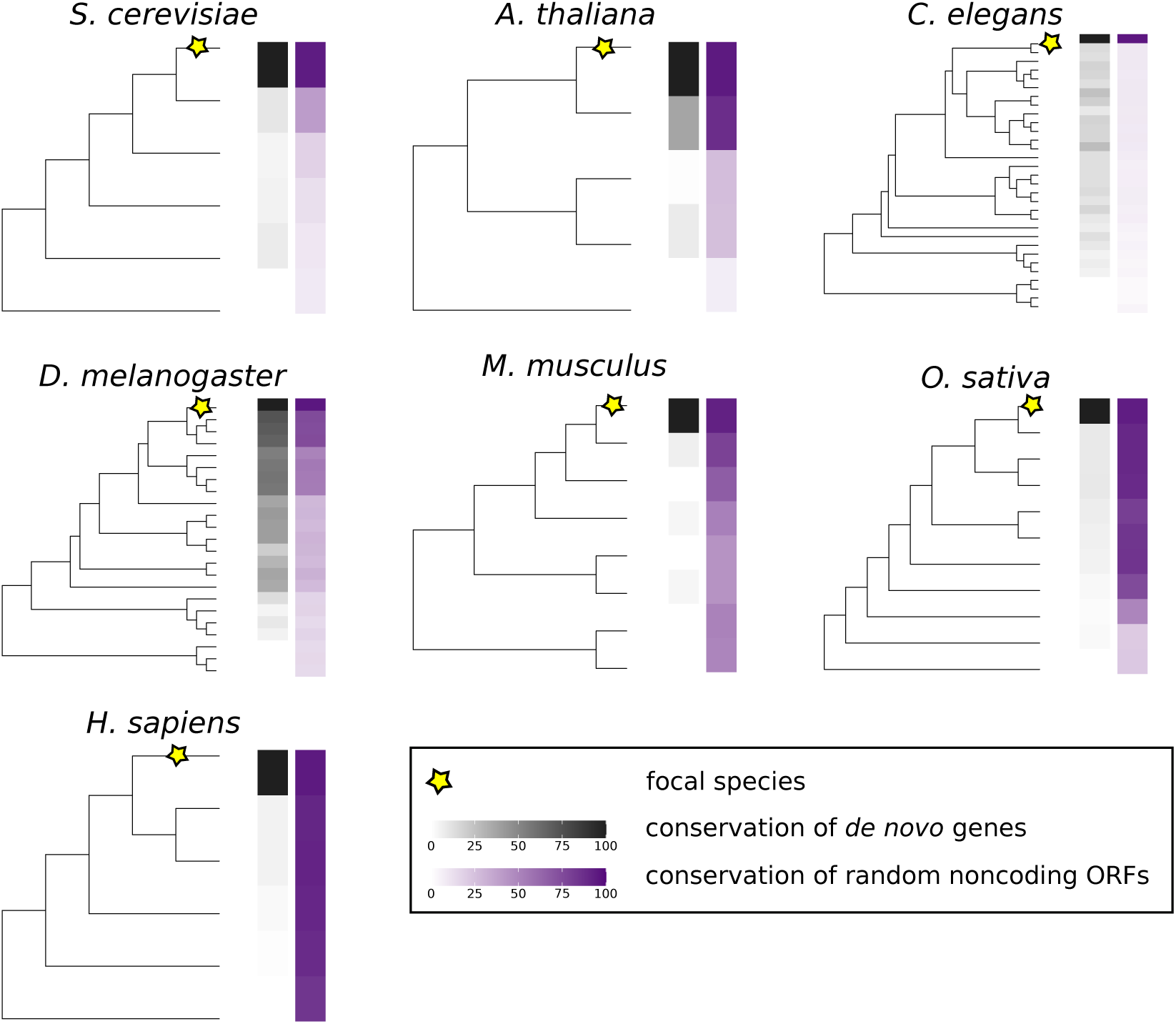
Conservation profiles of predicted de novo emerged genes and noncoding ORFs across the focal species’s neighbors. The trees are represented for the seven studied species and their neighbors (see Figure S1 for the trees with the names of species). Each tree is associated with two heatmaps. The left one (gradient of grays) represents the conservation levels of all the focal de novo emerged genes predicted by DENSE. The color of a given species in the tree represents the fraction of the focal de novo emerged genes that were detected until this species, with dark colors corresponding to species for which an important fraction of the focal de novo genes has been detected in it. The right heatmap (gradient of purple) represents the conservation level of a subset of 1000 noncoding ORFS selected randomly as calculated for the left heatmap.

Accordingly, we sought to compare for each focal species, our ability to detect the homology traces of its noncoding ORFs across its neighbors by establishing what we call a noncoding ORFs’ detectability profile. We therefore randomly selected subsets of 1000 noncoding ORFs from each focal genome and screened them with tblastn against the noncoding regions of the neighbors. We then calculated the fraction of these ORFs with a significant tblastn hit within the neighbors (Figure 3). This served as a proxy to estimate the speed at which the homology signal within neutrally evolving regions disappears among related species, providing us with a detectability profile that reflects the progression of noncoding ORFs’ detectability across the tree. The latter that is calculated between noncoding ORFs and noncoding regions gives a lower bounded estimation of the homology signal that can be expected between de novo emerged genes (i.e., under selection) and their orthologous noncoding regions (i.e., neutral evolution). For *S. cerevisiae, C. elegans, D. melanogaster, and A. thaliana,* the conservation of the predicted de novo emerged genes across the neighbors overall correlates with the detectability profiles of noncoding ORFs, suggesting that their detection is bounded by the detectability of the noncoding hits within the outgroup species (see Table S2 for the corresponding Spearman correlation coefficients). This opens the question of whether additional young de novo emerged genes are missed in these four species due to the high phylogenetic distance separating the focal and its closest neighbors. In contrast, in the case of *O. sativa,* as many as 75% of the randomly selected noncoding ORFs still exhibit significant hits until *O. meridionalis*, which may explain the higher count of de novo gene candidates predicted for this species (Figure 3, Figure S1). To assess the influence of the neighboring species proximity on the detection of de novo emerged genes, we iteratively designated each *O. sativa*‘s neighboring species as its new closest neighbor by systematically excluding neighbors that were phylogenetically closer than the newly assigned nearest neighbor (Figure 4A). Doing so, we effectively demonstrated that the prediction of de novo emerged genes in *O. sativa* is significantly impacted by the choice of the closest neighbor. The number of predicted de novo emerged genes diminishes rapidly with the distance to the closest considered neighbor. Notably, a sharp decrease in the number of predicted de novo emerged genes is observed when *O. punctata* is considered the nearest neighbor, aligning precisely with the lineage associated with a noticeable decline in the detectability of noncoding ORFs. This trend holds throughout the tree, as shown by the strong correlation between the number of predicted de novo emerged genes and the detectability profile of noncoding ORFs (Spearman’s correlation coefficient: Rho = –1, P = 1.7e-61). This result further strengthens the importance of the proximity of the neighbors considered in the prediction of de novo emerged genes, again questioning the hypothesis that other young de novo gene candidates may be missed for species whose neighbors are too distant. This also supports the utility of noncoding ORF detectability profiles as a valuable proxy for estimating our ability to identify de novo emerged genes.

**Fig. 4:**
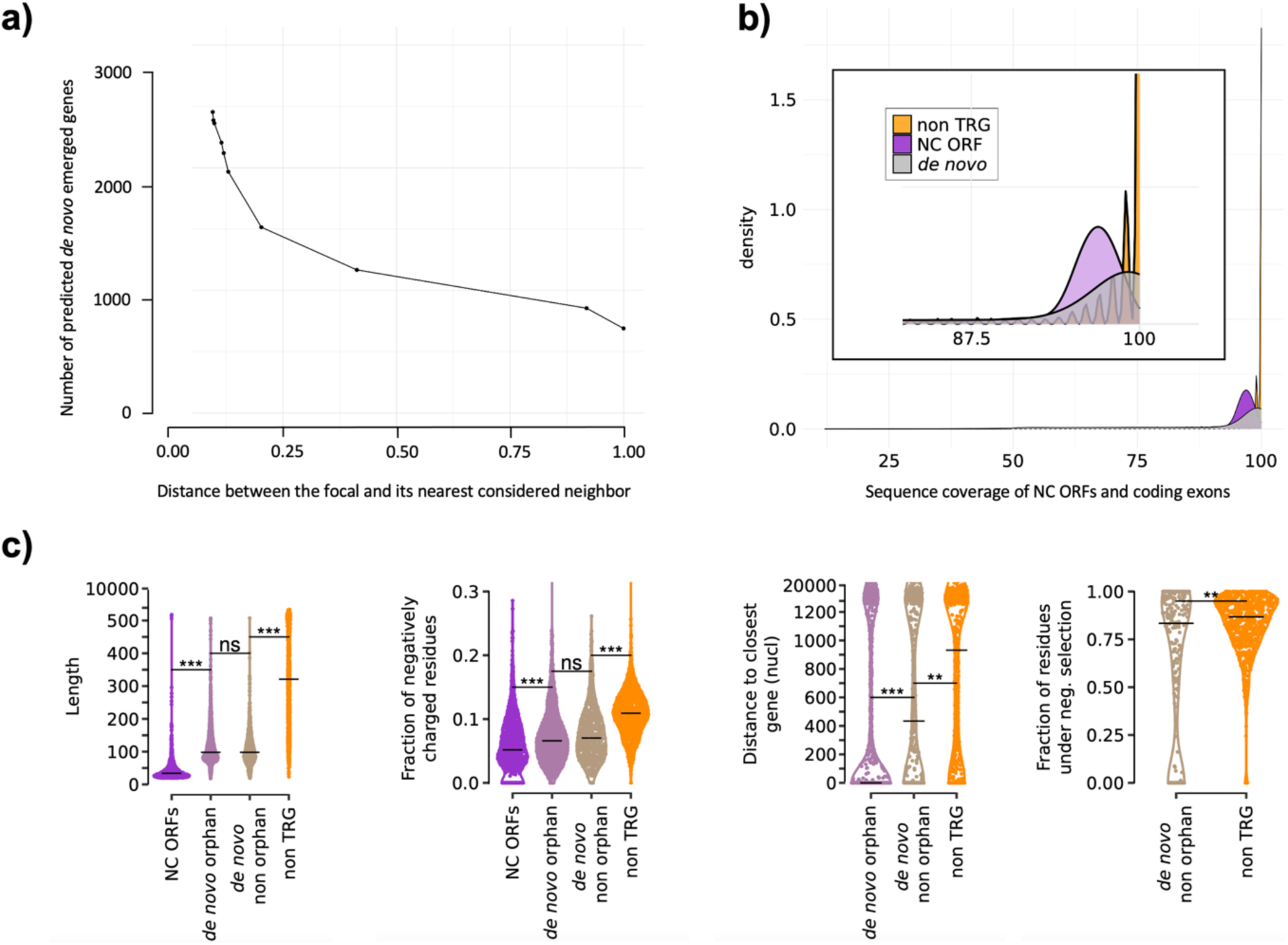
Analysis of the de novo emerged gene candidates of *O. sativa*. **a**) Number of de novo emerged genes that are predicted by DENSE (Strategy 1 with synteny criterion) with respect to the phylogenetic distance between the focal species and the nearest considered neighbor. The phylogenetic distance was calculated by OrthoFinder (Emms & Kelly, 2019). **b)** Distribution of the intactness of 1000 non-TRGs (orange), 1000 noncoding ORFs (purple), and the orphan de novo emerged genes (gray) of *O. sativa* across its neighboring species. The intactness represents the percentage of a noncoding ORF or coding sequence exon that is aligned with a homologous sequence detected in the neighbors (i.e., alignment coverage). **c)** Several properties calculated for 1000 noncoding ORFs and 1000 non-TRGs (purple and orange, respectively), and all the de novo genes of *O. sativa* separated in orphans and non-orphans. From left to right: ORF length (in amino acids), fraction of negatively charged residues, distance to the closest neighboring gene (in nucleotides), and fraction of residues undergoing negative selection. P-values were computed with the Mann–Whitney U test (one-sided). Asterisks denote level of significance: *, **, *** for P < 1×10−1, 1×10−2, 1×10−3, respectively.

Surprisingly, in the cases of *M. musculus* and *H. sapiens,* the number of predicted de novo emerged genes is very low despite a high noncoding ORFs’ detectability across their neighbors. In fact, the number of predicted de novo emerged genes is also limited by the count of detected TRGs, which is quite low for these two species compared to the others (see Table 1). However, whether this low number results from biological factors, from the phylostratum threshold chosen for these species (e.g., too young), or whether the higher numbers of TRGs detected in the other species simply reflect a potential taxonomic sampling bias remains unknown. Furthermore, in *O. sativa* and the two mammals, most of their de novo gene candidates (*O. sativa*: 77%, M. musculus 87%, *H. sapiens* 87%) are orphans, though, according to their noncoding ORFs’ detectability profiles, one might anticipate higher levels of de novo genes being shared by their closest neighbors. Yet, the fractions of de novo emerged genes conserved beyond the focal species remain very low (23%, 13%, and 13% for *O. sativa, M. musculus,* and *H. sapiens,* respectively). We can hypothesize that these genes emerged earlier and were subsequently lost in the sister lineages, recalling the high turnover of novel ORFs reported by Grandchamp et al. (Grandchamp et al., 2023); alternatively, they may reflect very young genes that have emerged recently in the lineage of the focal species, or they may simply result from annotation bias. Model organisms may have been over-annotated with respect to their neighbors as it is the case for example with the manual annotation of yeast available in the Saccharomyces Genome Database (Cherry et al., 2012). If so, the young genes detected in the focal species may also be present in the neighbors while not annotated as coding. However, one should note that for these three species, the genomes of the focal and the neighbors were annotated using the same pipeline (Ensembl 53 for *O. sativa* and NCBI for *M. musculus* and *H. sapiens*), indicating that the annotation bias, if any, are likely to be the same for all the compared genomes. In the following section, we chose to focus on the case of *O. sativa,* which harbors a significant number of de novo emerged genes for meaningful statistical analyses. To explore the possibility that the high number of de novo gene candidates specific to *O. sativa* could be attributed to potential over-annotation, we investigated whether their corresponding exons were intact in the neighboring species by calculating the proportion of each exon that was aligned with its homologous hits (i.e., exon coverage). A high level of exon intactness would suggest the presence of older genes that might have been overlooked when annotating the neighboring genomes. As a control, we repeated the experiment for a subset of 1000 non-TRGs (i.e., old CDSs) and 1000 noncoding ORFs randomly extracted from *O. sativa*. These subsets enabled us to estimate the conservation levels expected for established CDSs and noncoding ORFs, respectively, over this timescale. Firstly, it is worth noting that the exons of non-TRGs, on average, are retrieved in 85% of the screened *Oryza* species, while the noncoding ORFs and the exons of the orphan de novo emerged genes are less conserved, being detected in 70% of the genomes. Figure 4B represents the distribution of the alignment coverage of *O. sativa*’s noncoding ORFs and that of the exons of its non-TRGs and de novo genes when aligned with their homologs detected in the sister lineages (i.e., intactness). We show that the distribution of the intactness of de novo orphan exons lies between those of the noncoding ORFs and the non-TRG exons. On average, the coverage between the aligned homologous exons is significantly lower for de novo emerged genes than for older CDSs (Mann–Whitney U-test (one-sided), P = 1.6e-29), supporting the hypothesis that these young genes do not overall correspond to older genes missed by the annotation pipeline of the *O. sativa*‘s neighbors. As an additional control, we used DENSE to search for the de novo genes of the three closest neighbors of *O. sativa* which share the same topological position within the tree. Doing so, we detected comparable, albeit lower, numbers of de novo emerged genes (1451, 1762, and 2021 for *O. indica*, *O. nivara*, *O. rufipogon*, respectively), indicating that *Oryza* species, overall, exhibit higher numbers of de novo emerged genes. However, it remains difficult to disentangle the effective contribution of the close proximity of *Oryza* species from that of potential biological factors that could lead to high levels of de novo emerged genes. As observed for *O. sativa*, the majority of the de novo emerged genes of *O. indica* are orphans (59%)(Figure S2). *O. nivara* and *O. rufipogon* which are very close (about 700K years according to TimeTree (Kumar et al., 2022), share on average 44% of their de novo emerged genes and are respectively associated with 38% and 36% of orphan de novo genes. These results highlight that despite the close proximity of these species, the latter are nevertheless associated with high levels of de novo emerged genes specific to closely related lineages. Although the evolutionary fate of these genes is unpredictable, their remarkably low conservation levels suggest that a significant fraction of them may be short-lived in evolutionary history.

This prompted us to ask whether these ORFs are indeed coding and do not indicate noncoding ORFs erroneously annotated as coding. Determining whether an ORF is coding is a very difficult question. Nevertheless, the scientific community has identified a set of features that are typical hallmarks of coding sequences and that turned out to be efficient in characterizing young ORFs (Carvunis et al., 2012; Couso & Patraquim, 2017; Papadopoulos et al., 2021; Peng & Zhao, 2023). Figure 4C represents for *O. sativa*, several of these properties calculated for a subset of 1000 noncoding ORFs, 1000 non-TRGs, and all predicted de novo emerged genes classified as orphans or non-orphans. The ORF length has long been recognized as a key feature for detecting coding ORFs, with annotated CDSs being longer than noncoding ORFs regardless of the considered species. In *O. sativa*, genes predicted as de novo emerged, irrespective of being orphan or not, generally exhibit intermediate sizes between noncoding ORFs and old CDSs. It is important to note, however, that the ORF size is a feature that is explicitly taken into account in classical annotation pipelines, potentially introducing bias in the size of these young ORFs, since smaller ones may have been automatically excluded during the annotation process. Previously, we and others have demonstrated that CDSs in yeast and fly were enriched in negatively charged residues compared to noncoding ORFs, likely contributing to translation efficiency and/or preventing promiscuous interactions with the highly abundant and negatively charged ribosomes (Couso & Patraquim, 2017; Papadopoulos et al., 2021). This feature, therefore, offers a good proxy to interrogate the codability of subsets of candidates. Precisely, Figure 4C shows that the de novo gene candidates, including the orphan ones, display distributions of negatively charged residue fractions that lie between those of noncoding ORFs and old CDSs. This observation underscores that these candidates do not resemble noncoding ORFs and provides support for their classification as young recently de novo emerged genes. While not applicable to the orphan de novo gene candidates, the calculation of the dn/ds ratio for the non-orphan ones further substantiates this hypothesis with older de novo gene candidates exhibiting more than 62% of their residues under negative selection, a fraction comparable to that observed for older CDSs (see the Methods section for more details). Overall, the orphan de novo gene candidates resemble their non-orphan counterparts, except in their proximity to neighboring genes. Orphan candidates display significantly closer proximity to their surrounding genes than older de novo emerged genes and canonical CDSs. The genomic environment of nearby genes may contribute to their expression through transcription read-through, bidirectional promoter activity, or pervasive expression of regions of open chromatin (Gotea et al., 2013; McLysaght & Hurst, 2016; Wu & Sharp, 2013).

All these findings highlight the importance of having close neighbors in the detection of de novo emerged genes. Additionally, our analysis suggests that using very close species (i.e., associated with high noncoding ORFs’ detectability) may unveil populations of very young ORFs specific to the lineage of the focal species. Although the young ORFs specific to *O. sativa* overall resemble the older de novo gene candidates, the fact that most of them are not retrieved in the neighboring species (see Figure S2) suggests a rapid turnover of de novo emerged ORFs, at least in *Oryza* species, whose majority may be short-lived in evolutionary history, though the fate of each individual ORFs remains currently unpredictable.

### Application of DENSE to short timescales

In this section, we illustrate an application of DENSE to short timescales, which can be very useful when one aims to investigate the distribution and/or conservation of de novo emerged ORFs in a population. In fact, DENSE can handle genomes of different strains or lines, thereby enabling the characterization of the emergence of novel ORFs during very short timescales. In this example, we sought to assess the conservation of the de novo gene candidates identified in the reference line of *D. melanogaster*, in six other lines sequenced and annotated in Grandchamp et al. (Grandchamp et al., 2023). Therefore, we started from the TRGs of the reference line detected previously with the default parameters of DENSE (see Table 1), and directly entered the DENSE pipeline at the filtering step (Figure 1C). In this configuration, the focal is the reference line, and the neighbors consist of seven genomes of the *Drosophila* genus and those of the other six *D. melanogaster* lines (see the list of fly lines in Table S3). As in the previous section, genes are considered as de novo emerged according to the Strategy 1 combined with the synteny criterion. Figure 5A shows that all the de novo emerged genes predicted for the reference line are present in the other six fly lines. This result contrasts with the observation made in Grandchamp et al., where most de novo expressed ORFs (i.e., neoORFs that consist of not-yet fixed precursors of de novo genes) are generally observed in a single line, supporting a high birth-death rate (Figure 5B)(Grandchamp et al., 2023). Figure 5C represents the size and the fraction of negatively charged residues of these two ORF categories along with those of a subset of 1000 noncoding ORFs. The neoORFs and the de novo gene candidates display comparable size while being significantly longer than noncoding ORFs (Mann–Whitney U-test (one-sided), both P < 5.5×10-11). However, the fraction of negatively charged residues of the neoORFs is similar to that of noncoding ORFs and lower than that of the de novo emerged genes. This suggests two different populations, an older one consisting of ORFs conserved across *D. melanogaster* lines that are currently undergoing an optimization of their amino acid content, and a younger one that encompasses potential gene precursors that provide templates for de novo gene birth but are probably short-lived.

**Fig. 5:**
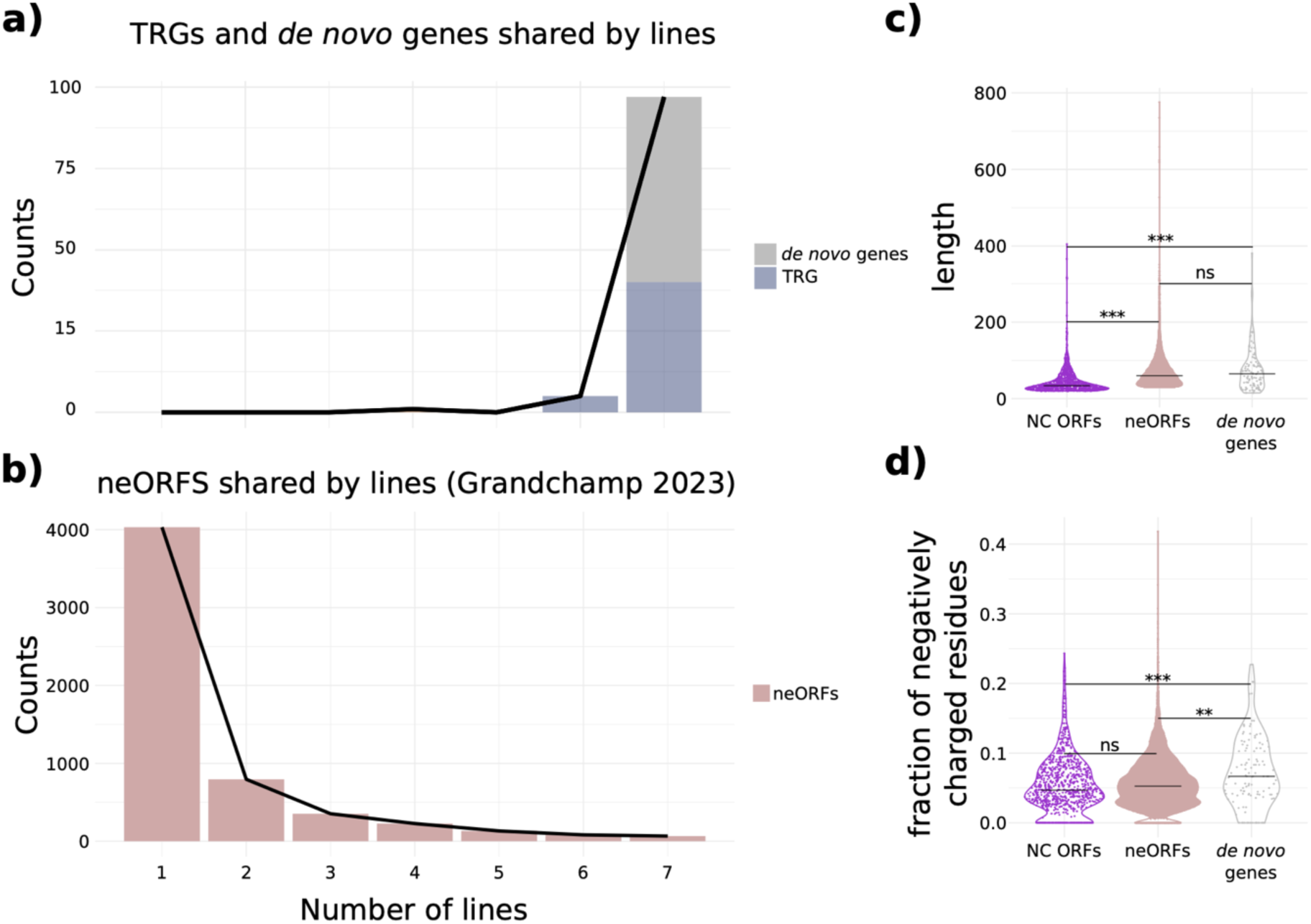
Conservation and properties of de novo emerged genes and neoORFs of *D. melanogaster*. **a**) Distribution of the conservation levels of the de novo emerged genes predicted for *D. melanogaster* with DENSE (Strategy 1 with synteny) across the seven fly lines. **b)** Same as panel a) for the neoORFs of *D. melanogaster* detected in Grandchamp et al., 2023. **c)** ORF length (top) and distributions of negatively charged residue fractions (bottom) for 1000 randomly selected noncoding ORFs of *D. melanogaster*, its neoORFs, and de novo emerged genes. Asterisks denote level of significance: *, **, *** for P < 1×10−1, 1×10−2, 1×10−3, respectively.

### Impact of parameters

We then investigated the impact of the different filters that can be applied to TRGs to classify them as de novo emerged. Unfortunately, no benchmark for de novo emerged genes is available, hampering the systematic study of parameter effects on the predictions. Furthermore, as de novo emerged genes get older, they become more challenging to detect since the criteria to classify them as de novo emerged (e.g., microsynteny, hit in noncoding regions of outgroup species) become more difficult to satisfy. As a result, older de novo emerged genes are likely to no longer conform to the constraints imposed by classical de novo gene detection protocols, and assessing whether individual candidates are true or false positives becomes a non-trivial task. Therefore, we deliberately removed some of the filters proposed by DENSE and analyzed the aforementioned properties of the resulting candidates, in order to see whether the latter are associated with properties similar to those of confidently identified candidates, i.e., with the complete set of filters as shown with the protocol presented in Figure 1 (Strategy 1 with synteny criterion).

Removing the synteny filter adds on average 67% of de novo emerged candidates for each of the seven focal species (see Table 1). Generally, despite slight significant differences, the candidates identified without synteny validation exhibit properties similar to those with a tblastn hit found in synteny, being overall not distinguishable according to their fractions of negatively charged residues or positions undergoing negative selection (Figure 6). In several species, they are, however, associated with longer size, higher distance to the next neighboring gene and higher phylogenetic distance (Figure 6). We may hypothesize that some of these candidates in fact correspond to de novo emerged genes, whose synteny with their orthologous noncoding region has been lost, and were consequently excluded with a stringent combination of filters. In fact, although the phylogenetic distance is not significantly higher for these candidates in four of the seven studied species, their conservation profiles reveal higher proportions of conserved de novo genes (Figure S1). This effect is even stronger for *O. sativa*, *H. sapiens,* and *C. elegans* which exhibit higher fractions of conserved de novo emerged genes when removing the synteny filter. This observation further supports the hypothesis of a population that includes slightly older de novo emerged genes whose genomic regions are no longer in synteny.

**Fig. 6:**
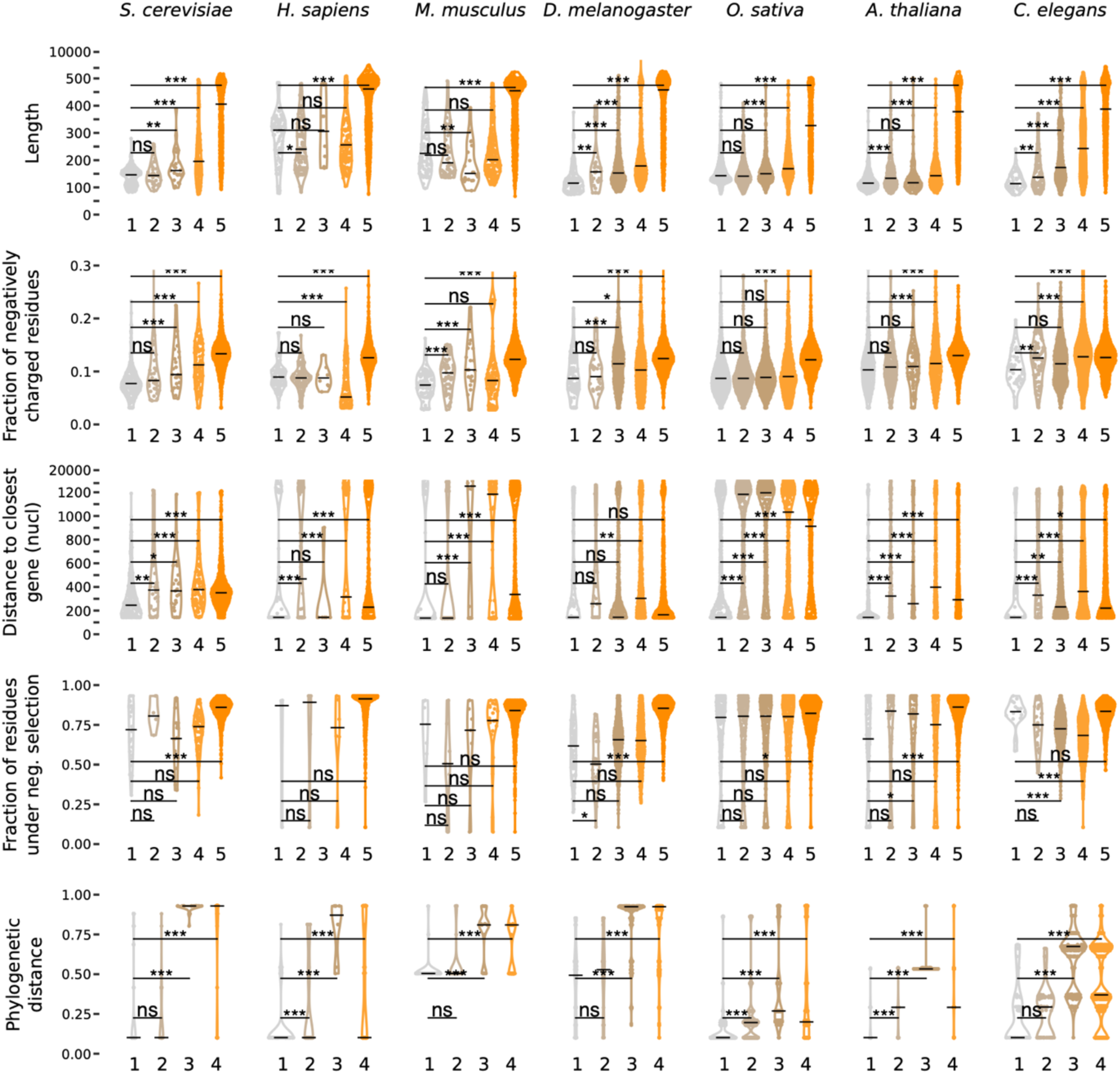
Properties of candidates detected with different combinations of criteria for each species. For all properties except Phylogenetic distance, **from left to right:** 1: de novo emerged gene predicted with a tblastn hit in a syntenic noncoding region of an outgroup species (strategy 1 with synteny), 2: de novo emerged gene candidates added when removing the synteny filter (strategy 1 without synteny), 3: de novo emerged gene candidates added when removing the requirement of the noncoding tblastn hit to be in an outgroup species (strategy 2 without synteny), 4: remaining TRGs, 5: and non-TRGs. Non-TRGs are absent from the Phylogenetic distance property. **From top to bottom:** Length of the corresponding CDSs (in amino acids). Fraction of negatively charged residues. Distance to the closest neighboring gene (in nucleotides). Fraction of positions under negative selection. Phylogenetic distance calculated with OrthoFinder between the focal species and the farthest species where the gene has been detected. P-values were computed with the Mann–Whitney U test (one-sided). Asterisks denote level of significance: *, **, *** for P < 1×10−1, 1×10−2, 1×10−3, respectively.

Candidates identified without the synteny criterion using Strategy 2 (i.e., those with a tblastn hit in the noncoding region of a neighboring species lacking the gene, irrespective of being outgroup or not, Figure 2A) display a more distinct separation from those identified with high confidence. For most species, they are associated with wider distributions than the de novo genes predicted with high confidence, unveiling a heterogeneous population of genes with various ages and probably different origins (Figure 6). These genes, identified with less stringent criteria, may include fast-evolving genes whose orthologs would have been lost in some neighboring species, explaining the detection of a tblastn hit in the noncoding regions of some neighbors. However, we do not exclude that it also comprises genes that emerged de novo million years ago, and for which homology in the noncoding orthologous regions of the outgroup species is no longer detectable. Finally, the ensemble of TRGs, as expected, consists of a highly heterogeneous population of genes with wider distributions for all considered features, reinforcing the idea that TRGs encompass genes with a wide diversity of origins and ages. It is interesting to note, that as filters are removed, the phylogenetic distance of the detected candidates increases, probably reflecting false positives with different ages, but also older de novo emerged genes that no longer satisfy the filter criteria.

Ultimately, we studied the influence of some parameters of the synteny filter on the number of validated de novo gene candidates. Specifically, we focused on two key factors: (i) the number of anchor pairs required in the vicinity of the tblastn hit, and (ii) the size of the window of genes that defined on each side of the focal de novo gene candidate and the noncoding hit in the outgroup species (Figure 2C, Figure 7). In Figure 7A, a candidate is classified as de novo emerged, if at least one anchor pair is found in the vicinity of its homologous noncoding hit of an outgroup species. This vicinity is defined by a given window size that varies from one to ten genes (see Figure 2C for more details). For all focal species, a window of four genes enables the detection of more than 84% of the candidates detected with the largest window (Figure 7A). This proportion significantly drops with smaller windows, reaching only 35% of the candidates detected with a window of one. Conversely, beyond four genes, the proportion of validated candidates increases gradually with the size of the window, suggesting that a window of four genes offers a suitable compromise. Figure 7B illustrates the impact of the number of anchor pairs, using a window of six, on the number of validated candidates. As expected, this number sharply diminishes with the number of anchor pairs, regardless of the focal species, reflecting the high speed at which the synteny is altered in related genomes.

**Fig. 7:**
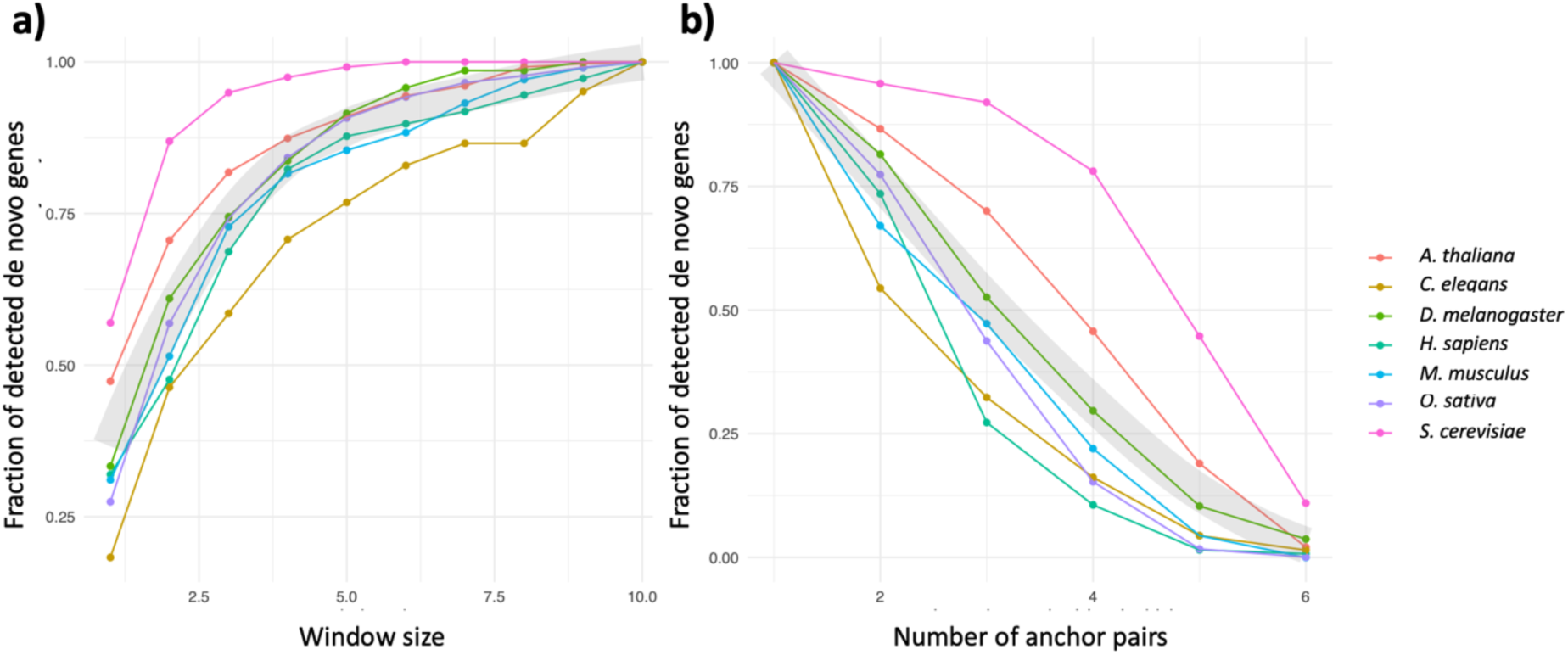
Impact of the window size and number of anchors used for the synteny search on the number of predicted de novo emerged genes. **a**) Fraction of predicted de novo emerged genes with respect to the size of the window (i.e., number of considered genes on each side of the focal de novo gene candidate). The fraction is calculated according to the number of de novo emerged genes predicted with a size window of 10. The number of anchor pairs (i.e., number of anchors per window) has been set to one. **b)** Fraction of predicted de novo emerged genes with respect to the number of required anchor pairs. The size of the window has been set to six. The fraction is calculated with respect to the number of de novo emerged genes predicted with a number of anchors set to one.

### DENSE online

The filtering part of the DENSE workflow, including all criteria combinations (Figure 1CD) is available through a web server once users provide their own list of TRGs (https://bioi2.i2bc.paris-saclay.fr/django/denovodb/dense-run/). Since, calculating the phylostratigraphy can be highly time-consuming, and as many studies involve the same model organisms, we also provide the DENSE calculations for the seven model organisms of the present study through a public database (https://bioi2.i2bc.paris-saclay.fr/django/denovodb/). These calculations include the phylostratigraphy calculated from the nr database (downloaded on March 23, 2022) and the predictions obtained from most of the DENSE available combinations of criteria.

## Discussion

In this study, we investigated the influence of different parameters in de novo emerged gene detection. Notably, we demonstrated the significant impact of the phylogenetic distance separating the focal species and its closest neighbors on the ability to detect de novo emerged genes, with the homology signal in noncoding regions decreasing rapidly as this distance increases. The decrease in the number of predicted candidates in *O. sativa* from 2455 to 1515, upon removing neighboring species with less than 1.5 million years of divergence, raises questions about the existence of a population of young candidates being missed in the other focal species. Additionally, we showed that the distance between the focal species and its nearest neighbors impacts the nature of the detected candidates. Indeed, the use of very close neighbors enables the detection of very young de novo emerged genes, such as the 1881 candidates specific to *O. sativa*. The latter, while displaying similar properties to the older de novo emerged genes, may not share the same fate. In particular, an important fraction of them might have a limited lifespan in evolutionary history, as suggested by the low amounts of de novo emerged genes shared across closely related neighbors. Furthermore, we found that removal of the synteny filter resulted in an average 67% increase in predicted de novo emerged genes in the seven studied species. These candidates, overall, do not differ significantly from those detected with a high degree of confidence. It can be therefore hypothesized that the latter include a population of de novo emerged genes whose synteny has been lost over time and which were subsequently excluded when the strictest combination of criteria was applied. Finally, omitting the requirement of a noncoding hit in an outgroup species led to a heterogeneous population of candidates with properties slightly different from those of candidates detected with high confidence. This population may encompass de novo emerged genes but also TRGs with other origins, underlining the difficulty of finding the set of parameters that minimizes the number of false positives and false negatives in the context of de novo emerged gene prediction.

In fact, for each species, there is a specific window of time during which de novo emerged gene candidates can be identified confidently. Beyond this, the signals used for reliable prediction are no longer detectable. As de novo emerged genes get older, their genetic context is likely to have changed relative to their orthologous noncoding region and homology traces within orthologous noncoding regions get blurred. As a result, determining the coding status of their ancestor becomes challenging, and classifying them as de novo emerged rapidly becomes impossible. However, although these de novo emerged genes are too old to be identified, they remain too young to resemble old, canonical CDSs, and overall exhibit intermediate properties between recently emerged de novo genes and canonical CDSs. While this behavior could be helpful in identifying them, it would require being also able to discriminate them from duplicated genes undergoing free-fall evolution, which, unfortunately, are also expected to be associated with intermediate properties (Montañés et al., 2023; Vakirlis et al., 2020). Predicting the coding or noncoding status of the ancestor is therefore key, and methods based on ancestral sequence reconstruction appear as a promising approach to distinguish emerging genes from those undergoing free-fall evolution. Nevertheless, the features typical of codability reported by the scientific community (Carvunis et al., 2012; Couso & Patraquim, 2017; McLysaght & Hurst, 2016; Papadopoulos et al., 2021; Peng & Zhao, 2023), while effective in characterizing a population of sequences, prove inadequate when dealing with a single sequence, again demonstrating the complexity of identifying de novo emerged genes outside a specific time window. The time window for the efficient detection of de novo emerged genes is not universal and determining the right one is not trivial. Indeed, geological time is not suitable for species comparison since evolution is not a linear process, species are associated with different generation times and have their own evolutionary trajectories. In other words, species have their own evolutionary time. In this work, we showed that characterizing the conservation profiles or intactness of both CDSs and noncoding ORFs across the tree offers a valuable route to delineating the boundaries of the evolutionary signal resulting from either selection (i.e., CDSs under selective pressure) or neutral evolution (i.e., noncoding ORFs). These profiles provide useful landmarks to estimate the upper and lower bounds of intact ORFs or detectable evolutionary traces expected in neighboring species associated with different divergence times, regardless of the knowledge of the generation time and evolutionary history of the considered species. Consequently, they can assist the user in adjusting the list of neighboring species or in identifying configurations where the nearest neighbors are too divergent for the accurate detection of de novo emerged genes.

It should be noted that the number of de novo emerged genes that can be detected is also bounded by the number of TRGs, since, by definition, the number of predicted de novo emerged genes cannot exceed the count of TRGs. This number is, in turn, directly affected by the taxonomic sampling around the focal species, the phylostratum used to define the TRGs, the heterogeneity in genome annotations across neighbors, and the sensitivity of the homology search when dating a species’ genome. Undersampling is likely to underestimate gene age by lacking the evolutionary relays that could connect them to their homologs in remote species, thus potentially leading to the misclassification of old genes as TRGs. In addition, the sensitivity of the homology search during the phylostratigraphy stage may impact the number of detected TRGs, as a lack of sensitivity can lead to underestimating gene age (Domazet-Lošo et al., 2017). On the other hand, heterogeneity in genome annotation among the compared species is also expected to lead to gene age underestimation by incorrectly categorizing genes that have been overlooked in other genomes as orphans. However, we showed that the characterization of the intactness of gene ORFs constitutes an efficient proxy to control this potential bias. Finally, choosing a phylostratum for the definition of TRGs that is too young inherently results in reduced lists of TRGs causing users to miss the genes whose emergence predates this phylostratum. If the genus level appears to be an effective phylostratum for most species in this study, the phylostratum threshold for *H. sapiens,* and *M. musculus* had to be adjusted to *Hominidae* and *Murinae,* respectively, to get sufficient numbers of outgroup species. Again, the conservation profiles of a subset of noncoding ORFs across the species associated with a given phylostratum can help users define the appropriate threshold. In any case, although defining the right phylostratum seems non-trivial in theory and directly depends on the species under consideration, in practice, false positives in TRG detection should be eliminated through the requirement of homology traces in the orthologous syntenic noncoding region of an outgroup species. This strict combination of criteria strongly supports the noncoding status of the ancestor. However, we do not exclude the possibility that a small fraction of genes that meet these criteria may consist of fast-evolving genes, which would have been recently lost in the sister lineages, thereby explaining the homology trace(s) detected in these species.

Finally, the main difficulty may stem from the term “de novo” genes itself which in fact, refers to the mechanism by which these genes have emerged. This semantic confusion, conflating the process with the product may inaccurately impart that de novo emerged genes are uniformly young and constitute a single and cohesive gene category, whereas, in reality, the population of de novo emerged genes is continuous and heterogeneous. It encompasses recently emerged genes but also, genes that appeared very early during evolution and whose origin is now unpredictable. This continuum implies a wide diversity of properties, functions, and trajectories, with recently emerged genes probably associated with uncertain fates, as suggested by the numerous young genes detected in *O. sativa*. In contrast, older de novo emerged genes have diversified over time, forging their unique trajectory during evolution. Consequently, all these genes, in all their diversity, share no more in common than their mechanism of emergence, and ultimately, their origin. Thus, which de novo emerged genes are we looking for? If the question is “How do we pass from the noncoding to the coding world”, part of the answer lies in the transition between these two worlds; specifically, in the study of genes that still bear the footprints of this transition. Precisely, the combination of criteria offered by DENSE, is tailored for the identification of these young genes that have recently emerged de novo.

## Conclusion

We introduced DENSE, a user-friendly Nextflow pipeline designed to seamlessly execute the entire protocol required for detecting de novo emerged genes from genomic data. This process encompasses the identification of TRGs through phylostratigraphy, along with their filtering according to various combinations of criteria. For higher specificity, we recommend employing the strictest protocol that relies on the filtering of candidates that exhibit homology traces in a syntenic noncoding region of an outgroup species. The latter, applicable genome-wide, stands out as the most promising for confirming the noncoding status of the ancestor using genomic data. It is important to note, however, that DENSE offers different combinations of strategies and parameters, empowering users to adapt to specific situations or explore new combinations.

The filtering step of DENSE is accessible to the scientific community through a web server, should users provide their own list of TRGs. Furthermore, as most studies focus on a limited set of model organisms, we have pre-calculated phylostratigraphies and executed the different DENSE strategies for the seven model organisms studied in this work. The associated results are available through a requestable database, that is to the best of our knowledge, the first public database of predicted de novo emerged genes. This unique dataset, encompassing seven model organisms and calculated with a consistent protocol, provides the scientific community with a valuable resource for cross-species analyses and large-scale studies. We plan to extend this database to other organisms and, hope that it will serve as a reference for de novo emerged gene lists generated with specific combinations of criteria. The integration of DENSE into a fully automated pipeline, and the modularity of its framework embedding different strategies and parameters should enable users to establish rational protocols for de novo emerged gene detection. This in turn should promote enhanced protocol communication, effective interoperability, and improved reproducibility across studies. While we anticipate that protocols will continue to evolve, we hope that this work, along with the rationality and interoperability facilitated by DENSE, will stimulate fruitful discussions and lead to further enhancements of protocols. Precisely, implemented through a Nextflow pipeline, DENSE is perfectly suited to these collaborative goals.

## Materials and Methods

### Identification of the de novo emerged genes in the seven studied organisms

All de novo emerged genes were predicted using the full pipeline of DENSE with default parameters (Strategy 1 with synteny, one anchor pair, and a window size of 4). For TRG detection, we used the nr database downloaded on March 23, 2022. The phylostratum threshold used for TRG detection was set to the genus level, except for *M. musculus* and *H. sapiens*. For these two latter, to ensure a sufficient number of outgroup species, the threshold was extended to *Murinae* and *Hominidae*, respectively. It is worth noting that the neighboring genomes used for detecting coding and noncoding hits during the TRG filtering process (Figure 1CD) may not correspond to all genomes included in the phylostratum selected for TRG detection. Notably, only species belonging to the *D. melanogaster* subgroup were considered for the filtering step. The list of all genomes used in this study, along with the links to their sequence and annotation files, are available in Supplementary Table S1. For each focal species, the associated local trees (focal and neighboring species) were generated using OrthoFinder (Emms & Kelly, 2019) with default parameters, except for the ‘msa’ method that was used for gene tree inference (‘-M’ option).

### ORF properties

Except for the calculation of distance to the closest neighboring gene, all studied properties were computed from the CDSs of the genes, including the de novo genes, orphan de novo genes, non-orphan de novo genes, TRGs, and non-TRGs. In cases where genes were associated with multiple isoforms, the evaluated properties were calculated on all their corresponding CDSs. The phylogenetic distance was directly extracted from the tree computed by OrthoFinder (Emms & Kelly, 2019). The fraction of residues under negative selection was calculated with codeml (Yang, 2007). Therefore, for each considered gene in the focal species, we searched for its orthologs within the neighboring species using the Best Reciprocal Hits method (E-value: 10-3, coverage: 70%). The corresponding CDSs were aligned with MAFFT (Katoh & Standley, 2013) and provided to codeml. We employed the model assuming a fixed Ω value along the branches and three states per site (negative, neutral, and positive). The distance to the closest neighboring gene was calculated using BedTools (Quinlan & Hall, 2010).

### Homology detection

The conservation of the de novo genes across the neighboring species (Figure 3), and the intactness of the three ORF categories (Figure 4B), were assessed using blastp (E-value: 10-3, coverage: 50%). The search for the homology signal of the focal noncoding ORFs in the neighboring species was realized using tblastn (E-value: 10-3, coverage: 50%).

### Statistical analyses

All statistical tests were performed in R (4.3.2) (R Core Team, 2021). When samples were larger than 500 individuals, tests were performed 100 times on random subsets of 500 individuals chosen from the initial sample, and the averaged P-value was subsequently calculated.

## Data Availability Statement

The complete list of genomes used for this study is provided in the Supplementary Table S1.

## Competing interest

The authors declare that they have no competing interests.

## Acknowledgments

We thank those associated with the Caenorhabditis Genomes Project for pre-publication access to genome data. We thank Ambre Baumann and Simon Herman for their assistance in testing the DENSE pipeline. PR’s work was supported by a French government fellowship. AG’s work was supported by the Deutsche Forschungsgemeinschaft priority program ‘Genomic Basis of Evolutionary Innovations’ (SPP 2349) BO 2544/20-1.

## Authors’ contributions

Conceptualization: PR, AL; Development and Investigations: PR, AG, CQ, AL; Writing: PR, AG, AL; Supervision: AL.

## Notes

### Competing Interest Statement

The authors have declared no competing interest.

