## Supplemental Information for "DE Novo emerged gene SEarch in Eukaryotes with DENSE"

### Tables of Contents

**Supplemental Figure S1:** Conservation profiles of predicted *de novo* emerged genes and noncoding ORFs across the focal species's neighbors.

**Supplemental Figure S2:** Conservation profiles of predicted *de novo* emerged genes in *O. sativa* and its three closest neighbors.

**Supplemental Table S1:** Organisms' meta data.

**Supplemental Table S2:** Correlation between the conservation of the *de novo* emerged genes and the conservation of noncoding ORFs across the neighbor species.

**Supplemental Table S3:** *D. melanogaster* lines.

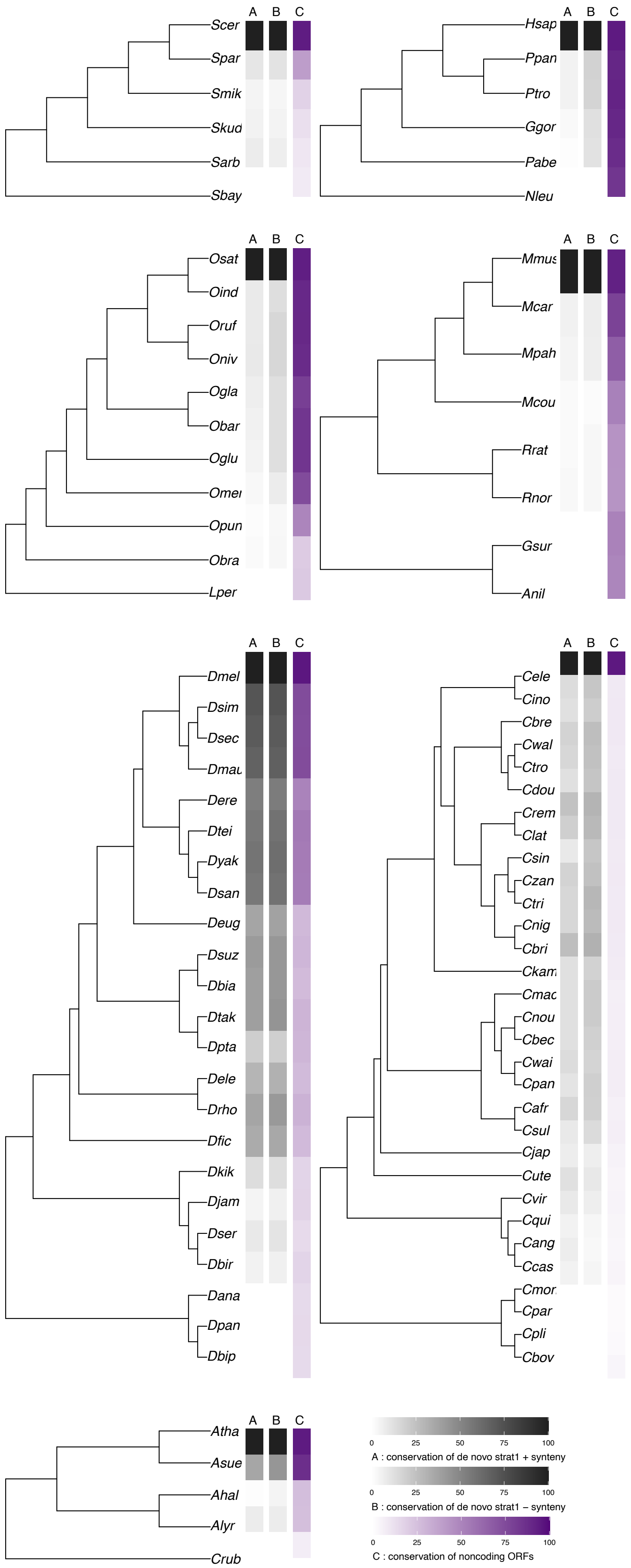
A B C
Scer
Spar
Smik
Skud
Sarb
Sbay
Hsap
Ppan
Ptro
Ggor
Pabe
Nleu
Osat
Oind
Oruf
Oniv
Ogla
Obar
Oglu
Omer
Opun
Obra
Lper
Mmus
Mcar
Mpah
Mcou
Rrat
Rnor
Gsur
Anil
Dmel
Dsim
Dsec
Dmau
Dere
Dtei
Dyak
Dsan
Deug
Dsuz
Dbia
Dtak
Dpta
Dele
Drho
Dfic
Dkik
Djam
Dser
Dbir
Dana
Dpan
Dbip
Cele
Cino
Cbre
Cwal
Ctro
Cdou
Crem
Clat
Csin
Czan
Ctri
Cnig
Cbri
Ckam
Cmac
Cnou
Cbec
Cwai
Cpan
Cafr
Csul
Cjap
Cute
Cvir
Cqui
Cang
Ccas
Cmor
Cpar
Cpli
Cbov
Atha
Asue
Ahal
Alyr
Crub
0 25 50 75 100
A : conservation of de novo strat1 + synteny
0 25 50 75 100
B : conservation of de novo strat1 – synteny
0 25 50 75 100
C : conservation of noncoding ORFs

**Supplemental Figure 1. Conservation profiles of predicted *de novo* emerged genes and noncoding ORFs across the focal species's neighbors.** Same color scheme as Figure 3. The second heatmap (B) (Strategy 1 only) represents the conservation of the *de novo* emerged genes predicted without synteny. The two other heatmaps (A and C) are the same as in Figure 3.

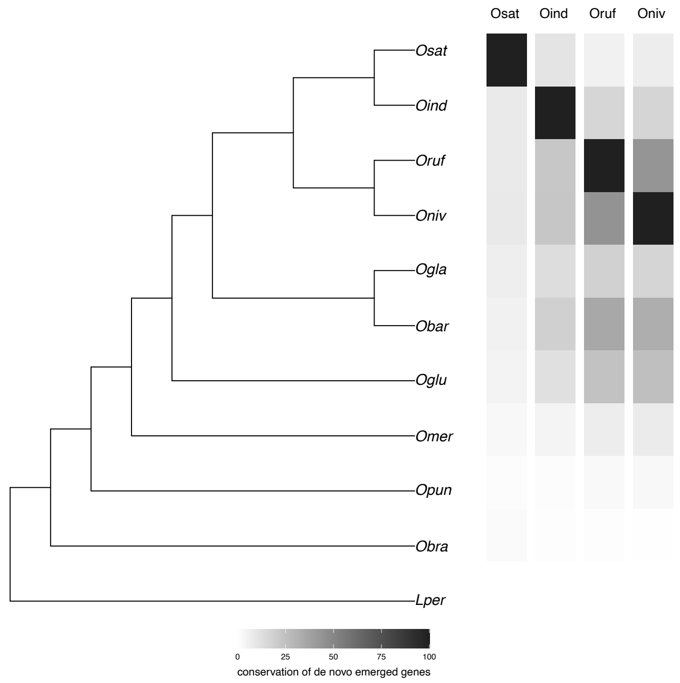


**Supplemental Figure 2. Conservation profiles of predicted de novo emerged genes in O. sativa and its three closest neighbors.** Same color scheme as Figure 3. The first one shows the conservation of the *de novo* genes predicted for *O. sativa* (focal species) across the tree (same heatmap as in Figure 4), while the others represent the conservation of the predicted de novo emerged genes of *O. indica*, *O. rufipogon*, and *O. nivara* respectively.

| organism | short name | taxid | genomic FASTA | GFF3 |
|---|---|---|---|---|
| *Saccharomyces cerevisiae* | | | | |
| *Saccharomyces arboricola* | Sarb | 706196 | http://bim.i2bc.paris-saclay.fr/anne-lopes/genomes_Roginski_GBE/Sacc_genomes_nonNCBI.tar.gz | http://bim.i2bc.paris-saclay.fr/anne-lopes/genomes_Roginski_GBE/Sacc_genomes_nonNCBI.tar.gz |
| *Saccharomyces bayanus* | Sbay | 4931 | http://bim.i2bc.paris-saclay.fr/anne-lopes/genomes_Roginski_GBE/Sacc_genomes_nonNCBI.tar.gz | http://bim.i2bc.paris-saclay.fr/anne-lopes/genomes_Roginski_GBE/Sacc_genomes_nonNCBI.tar.gz |
| *Saccharomyces cerevisiae S288C* | Scer | 559292 | https://www.ncbi.nlm.nih.gov/datasets/genome/GCF_000146045.2 | https://www.ncbi.nlm.nih.gov/datasets/genome/GCF_000146045.2 |
| *Saccharomyces kudriavzevii* | Skud | 114524 | http://bim.i2bc.paris-saclay.fr/anne-lopes/genomes_Roginski_GBE/Sacc_genomes_nonNCBI.tar.gz | http://bim.i2bc.paris-saclay.fr/anne-lopes/genomes_Roginski_GBE/Sacc_genomes_nonNCBI.tar.gz |
| *Saccharomyces mikatae* | Smik | 114525 | http://bim.i2bc.paris-saclay.fr/anne-lopes/genomes_Roginski_GBE/Sacc_genomes_nonNCBI.tar.gz | http://bim.i2bc.paris-saclay.fr/anne-lopes/genomes_Roginski_GBE/Sacc_genomes_nonNCBI.tar.gz |
| *Saccharomyces paradoxus* | Spar | 27291 | https://www.ncbi.nlm.nih.gov/datasets/genome/GCF_002079055.1 | https://www.ncbi.nlm.nih.gov/datasets/genome/GCF_002079055.1 |
| *Homo sapiens* | | | | |
| *Gorilla gorilla* | Ggor | 9593 | https://www.ncbi.nlm.nih.gov/datasets/genome/GCF_008122165.1 | https://www.ncbi.nlm.nih.gov/datasets/genome/GCF_008122165.1 |
| *Homo sapiens* | Hsap | 9606 | https://www.ncbi.nlm.nih.gov/datasets/genome/GCF_000001405.40 | https://www.ncbi.nlm.nih.gov/datasets/genome/GCF_000001405.40 |
| *Nomascus leucogenys* | Nleu | 61853 | https://www.ncbi.nlm.nih.gov/datasets/genome/GCF_006542625.1 | https://www.ncbi.nlm.nih.gov/datasets/genome/GCF_006542625.1 |
| *Pongo abelii* | Pabe | 9601 | https://www.ncbi.nlm.nih.gov/datasets/genome/GCF_002880775.1 | https://www.ncbi.nlm.nih.gov/datasets/genome/GCF_002880775.1 |
| *Pan paniscus* | Ppan | 9597 | https://www.ncbi.nlm.nih.gov/datasets/genome/GCF_013052645.1 | https://www.ncbi.nlm.nih.gov/datasets/genome/GCF_013052645.1 |
| *Pan troglodytes* | Ptro | 9598 | https://www.ncbi.nlm.nih.gov/datasets/genome/GCF_002880755.1 | https://www.ncbi.nlm.nih.gov/datasets/genome/GCF_002880755.1 |
| *Mus musculus* | | | | |
| *Arvicanthis niloticus* | Anil | 61156 | https://www.ncbi.nlm.nih.gov/datasets/genome/GCF_011762505.1 | https://www.ncbi.nlm.nih.gov/datasets/genome/GCF_011762505.1 |
| *Grammomys surdaster* | Gsur | 491861 | https://www.ncbi.nlm.nih.gov/datasets/genome/GCF_004785775.1 | https://www.ncbi.nlm.nih.gov/datasets/genome/GCF_004785775.1 |
| *Mus caroli* | Mcar | 10089 | https://www.ncbi.nlm.nih.gov/datasets/genome/GCF_900094665.1 | https://www.ncbi.nlm.nih.gov/datasets/genome/GCF_900094665.1 |
| *Mastomys coucha* | Mcou | 35658 | https://www.ncbi.nlm.nih.gov/datasets/genome/GCF_008632895.1 | https://www.ncbi.nlm.nih.gov/datasets/genome/GCF_008632895.1 |
| *Mus musculus* | Mmus | 10090 | https://www.ncbi.nlm.nih.gov/datasets/genome/GCF_000001635.27 | https://www.ncbi.nlm.nih.gov/datasets/genome/GCF_000001635.27 |
| *Mus pahari* | Mpah | 10093 | https://www.ncbi.nlm.nih.gov/datasets/genome/GCF_900095145.1 | https://www.ncbi.nlm.nih.gov/datasets/genome/GCF_900095145.1 |
| *Rattus norvegicus* | Rnor | 10116 | https://www.ncbi.nlm.nih.gov/datasets/genome/GCF_015227675.2 | https://www.ncbi.nlm.nih.gov/datasets/genome/GCF_015227675.2 |
| *Rattus rattus* | Rrat | 10117 | https://www.ncbi.nlm.nih.gov/datasets/genome/GCF_011064425.1 | https://www.ncbi.nlm.nih.gov/datasets/genome/GCF_011064425.1 |
| *Drosophila melanogaster* | | | | |
| *Drosophila erecta* | Dere | 7220 | https://www.ncbi.nlm.nih.gov/datasets/genome/GCF_003286155.1 | https://www.ncbi.nlm.nih.gov/datasets/genome/GCF_003286155.1 |
| *Drosophila eugracilis* | Deug | 29029 | https://www.ncbi.nlm.nih.gov/datasets/genome/GCF_018153835.1 | https://www.ncbi.nlm.nih.gov/datasets/genome/GCF_018153835.1 |
| *Drosophila mauritiana* | Dmau | 7226 | https://www.ncbi.nlm.nih.gov/datasets/genome/GCF_004382145.1 | https://www.ncbi.nlm.nih.gov/datasets/genome/GCF_004382145.1 |
| *Drosophila melanogaster* | Dmel | 7227 | https://www.ncbi.nlm.nih.gov/datasets/genome/GCF_000001215.4 | https://www.ncbi.nlm.nih.gov/datasets/genome/GCF_000001215.4 |
| *Drosophila ananassae* | Dana | 7217 | https://www.ncbi.nlm.nih.gov/datasets/genome/GCF_017639315.1 | https://www.ncbi.nlm.nih.gov/datasets/genome/GCF_017639315.1 |
| *Drosophila biarmipes* | Dbia | 125945 | https://www.ncbi.nlm.nih.gov/datasets/genome/GCF_025231255.1 | https://www.ncbi.nlm.nih.gov/datasets/genome/GCF_025231255.1 |
| *Drosophila bipectinata* | Dbip | 42026 | https://www.ncbi.nlm.nih.gov/datasets/genome/GCF_018153845.1 | https://www.ncbi.nlm.nih.gov/datasets/genome/GCF_018153845.1 |
| *Drosophila birchii* | Dbir | 46829 | https://www.ncbi.nlm.nih.gov/datasets/genome/GCA_021223725.1 | https://www.ncbi.nlm.nih.gov/datasets/genome/GCA_021223725.1 |
| *Drosophila elegans* | Dele | 30023 | https://www.ncbi.nlm.nih.gov/datasets/genome/GCF_018152505.1 | https://www.ncbi.nlm.nih.gov/datasets/genome/GCF_018152505.1 |
| *Drosophila ficusphila* | Dfic | 30025 | https://www.ncbi.nlm.nih.gov/datasets/genome/GCF_018152265.1 | https://www.ncbi.nlm.nih.gov/datasets/genome/GCF_018152265.1 |
| *Drosophila jambulina* | Djam | 111875 | https://www.ncbi.nlm.nih.gov/datasets/genome/GCA_021223805.1 | https://www.ncbi.nlm.nih.gov/datasets/genome/GCA_021223805.1 |
| *Drosophila kikkawai* | Dkik | 30033 | https://www.ncbi.nlm.nih.gov/datasets/genome/GCF_018152535.1 | https://www.ncbi.nlm.nih.gov/datasets/genome/GCF_018152535.1 |
| *Drosophila pseudoananassae* | Dpan | 65964 | https://www.ncbi.nlm.nih.gov/datasets/genome/GCA_021223845.1 | https://www.ncbi.nlm.nih.gov/datasets/genome/GCA_021223845.1 |
| *Drosophila pseudotakahashii* | Dpta | 193234 | https://www.ncbi.nlm.nih.gov/datasets/genome/GCA_021223935.1 | https://www.ncbi.nlm.nih.gov/datasets/genome/GCA_021223935.1 |
| *Drosophila rhopaloa* | Drho | 1041015 | https://www.ncbi.nlm.nih.gov/datasets/genome/GCF_018152115.1 | https://www.ncbi.nlm.nih.gov/datasets/genome/GCF_018152115.1 |
| *Drosophila serrata* | Dser | 7274 | https://www.ncbi.nlm.nih.gov/datasets/genome/GCF_002093755.2 | https://www.ncbi.nlm.nih.gov/datasets/genome/GCF_002093755.2 |
| *Drosophila suzukii* | Dsuz | 28584 | https://www.ncbi.nlm.nih.gov/datasets/genome/GCF_013340165.1 | https://www.ncbi.nlm.nih.gov/datasets/genome/GCF_013340165.1 |
| *Drosophila takahashii* | Dtak | 29030 | https://www.ncbi.nlm.nih.gov/datasets/genome/GCF_018152695.1 | https://www.ncbi.nlm.nih.gov/datasets/genome/GCF_018152695.1 |
| *Drosophila santomea* | Dsan | 129105 | https://www.ncbi.nlm.nih.gov/datasets/genome/GCF_016746245.2 | https://www.ncbi.nlm.nih.gov/datasets/genome/GCF_016746245.2 |
| *Drosophila sechellia* | Dsec | 7238 | https://www.ncbi.nlm.nih.gov/datasets/genome/GCF_004382195.2 | https://www.ncbi.nlm.nih.gov/datasets/genome/GCF_004382195.2 |
| *Drosophila simulans* | Dsim | 7240 | https://www.ncbi.nlm.nih.gov/datasets/genome/GCF_016746395.2 | https://www.ncbi.nlm.nih.gov/datasets/genome/GCF_016746395.2 |
| *Drosophila teissieri* | Dtei | 7243 | https://www.ncbi.nlm.nih.gov/datasets/genome/GCF_016746235.2 | https://www.ncbi.nlm.nih.gov/datasets/genome/GCF_016746235.2 |
| *Drosophila yakuba* | Dyak | 7245 | https://www.ncbi.nlm.nih.gov/datasets/genome/GCF_016746365.2 | https://www.ncbi.nlm.nih.gov/datasets/genome/GCF_016746365.2 |
| *Oryza sativa* | | | | |
| *Leersia perrieri* | Lper | 77586 | https://ftp.ebi.ac.uk/ensemblgenomes/pub/release-53/plants/fasta/leersia_perrieri/dna/Leersia_perrieri.Lperr_V1.4.dna.toplevel.fa.gz | https://ftp.ebi.ac.uk/ensemblgenomes/pub/release-53/plants/gff3/leersia_perrieri/Leersia_perrieri.Lperr_V1.4.53.gff3.gz |
| *Oryza barthii* | Obar | 65489 | https://ftp.ebi.ac.uk/ensemblgenomes/pub/release-53/plants/fasta/oryza_barthii/dna/Oryza_barthii.O.barthii_v1.dna.toplevel.fa.gz | https://ftp.ebi.ac.uk/ensemblgenomes/pub/release-53/plants/gff3/oryza_barthii/Oryza_barthii.O.barthii_v1.53.gff3.gz |
| *Oryza brachyantha* | Obra | 4533 | https://ftp.ebi.ac.uk/ensemblgenomes/pub/release-53/plants/fasta/oryza_brachyantha/dna/Oryza_brachyantha.Oryza_brachyantha.v1.4b.dna.toplevel.fa.gz | https://ftp.ebi.ac.uk/ensemblgenomes/pub/release-53/plants/gff3/oryza_brachyantha/Oryza_brachyantha.Oryza_brachyantha.v1.4b.53.gff3.gz |
| *Oryza glaberrima* | Ogla | 4538 | https://ftp.ebi.ac.uk/ensemblgenomes/pub/release-53/plants/fasta/oryza_glaberrima/dna/Oryza_glaberrima.Oryza_glaberrima_V1.dna.toplevel.fa.gz | https://ftp.ebi.ac.uk/ensemblgenomes/pub/release-53/plants/gff3/oryza_glaberrima/Oryza_glaberrima.Oryza_glaberrima_V1.53.gff3.gz |
| *Oryza glumipatula* | Oglu | 40148 | https://ftp.ebi.ac.uk/ensemblgenomes/pub/release-53/plants/fasta/oryza_glumipatula/dna/Oryza_glumipatula.Oryza_glumaepatula_v1.5.dna.toplevel.fa.gz | https://ftp.ebi.ac.uk/ensemblgenomes/pub/release-53/plants/gff3/oryza_glumipatula/Oryza_glumipatula.Oryza_glumaepatula_v1.5.53.gff3.gz |
| *Oryza sativa Indica Group* | Oind | 39946 | https://ftp.ebi.ac.uk/ensemblgenomes/pub/release-53/plants/fasta/oryza_indica/dna/Oryza_indica.ASM465v1.dna.toplevel.fa.gz | https://ftp.ebi.ac.uk/ensemblgenomes/pub/release-53/plants/gff3/oryza_indica/Oryza_indica.ASM465v1.53.gff3.gz |
| *Oryza meridionalis* | Omer | 40149 | https://ftp.ebi.ac.uk/ensemblgenomes/pub/release-53/plants/fasta/oryza_meridionalis/dna/Oryza_meridionalis.Oryza_meridionalis_v1.3.dna.toplevel.fa.gz | https://ftp.ebi.ac.uk/ensemblgenomes/pub/release-53/plants/gff3/oryza_meridionalis/Oryza_meridionalis.Oryza_meridionalis_v1.3.53.gff3.gz |
| *Oryza nivara* | Oniv | 4536 | https://ftp.ebi.ac.uk/ensemblgenomes/pub/release-53/plants/fasta/oryza_nivara/dna/Oryza_nivara.Oryza_nivara_v1.0.dna.toplevel.fa.gz | https://ftp.ebi.ac.uk/ensemblgenomes/pub/release-53/plants/gff3/oryza_nivara/Oryza_nivara.Oryza_nivara_v1.0.53.gff3.gz |
| *Oryza punctata* | Opun | 4537 | https://ftp.ebi.ac.uk/ensemblgenomes/pub/release-53/plants/fasta/oryza_punctata/dna/Oryza_punctata.Oryza_punctata_v1.2.dna.toplevel.fa.gz | https://ftp.ebi.ac.uk/ensemblgenomes/pub/release-53/plants/gff3/oryza_punctata/Oryza_punctata.Oryza_punctata_v1.2.53.gff3.gz |
| *Oryza rufipogon* | Oruf | 4529 | https://ftp.ebi.ac.uk/ensemblgenomes/pub/release-53/plants/fasta/oryza_rufipogon/dna/Oryza_rufipogon.OR_W1943.dna.toplevel.fa.gz | https://ftp.ebi.ac.uk/ensemblgenomes/pub/release-53/plants/gff3/oryza_rufipogon/Oryza_rufipogon.OR_W1943.53.gff3.gz |
| *Oryza sativa Japonica Group* | Osat | 39947 | https://ftp.ebi.ac.uk/ensemblgenomes/pub/release-53/plants/fasta/oryza_sativa/dna/Oryza_sativa.IRGSP-1.0.dna.toplevel.fa.gz | https://ftp.ebi.ac.uk/ensemblgenomes/pub/release-53/plants/gff3/oryza_sativa/Oryza_sativa.IRGSP-1.0.53.gff3.gz |
| *Arabidopsis thaliana* | | | | |
| *Arabidopsis halleri subsp. gemmifera* | Ahal | 63677 | https://ftp.ebi.ac.uk/ensemblgenomes/pub/release-53/plants/fasta/arabidopsis_halleri/dna/Arabidopsis_halleri.Ahal2.2.dna.toplevel.fa.gz | https://ftp.ebi.ac.uk/ensemblgenomes/pub/release-53/plants/gff3/arabidopsis_halleri/Arabidopsis_halleri.Ahal2.2.53.gff3.gz |
| *Arabidopsis lyrata subsp. lyrata* | Alyr | 81972 | https://www.ncbi.nlm.nih.gov/datasets/genome/GCF_000004255.2 | https://www.ncbi.nlm.nih.gov/datasets/genome/GCF_000004255.2 |
| *Arabidopsis suecica* | Asue | 45249 | https://www.ncbi.nlm.nih.gov/datasets/genome/GCA_019202805.1 | https://www.ncbi.nlm.nih.gov/datasets/genome/GCA_019202805.1 |
| *Arabidopsis thaliana* | Atha | 3702 | https://www.ncbi.nlm.nih.gov/datasets/genome/GCF_000001735.4 | https://www.ncbi.nlm.nih.gov/datasets/genome/GCF_000001735.4 |

| | | | | |
|---|---|---|---|---|
| *Capsella rubella* | Crub | 81985 | https://www.ncbi.nlm.nih.gov/datasets/genome/GCF_000375325.1 | https://www.ncbi.nlm.nih.gov/datasets/genome/GCF_000375325.1 |
| | | *Caenorhabditis elegans* | | |
| *Caenorhabditis angaria* | Cang | 860376 | https://ftp.ebi.ac.uk/pub/databases/wormbase/parasite/releases/WBPS16/species/caenorhabditis_angaria/PRJNA51225/caenorhabditis_angaria.PRJNA51225.WBPS16.genomic.fa.gz | https://ftp.ebi.ac.uk/pub/databases/wormbase/parasite/releases/WBPS16/species/caenorhabditis_angaria/PRJNA51225/caenorhabditis_angaria.PRJNA51225.WBPS16.annotations.gff3.gz |
| *Caenorhabditis becei* | Cbec | 2301260 | https://ftp.ebi.ac.uk/pub/databases/wormbase/parasite/releases/WBPS16/species/caenorhabditis_becei/PRJEB28243/caenorhabditis_becei.PRJEB28243.WBPS16.genomic.fa.gz | https://ftp.ebi.ac.uk/pub/databases/wormbase/parasite/releases/WBPS16/species/caenorhabditis_becei/PRJEB28243/caenorhabditis_becei.PRJEB28243.WBPS16.annotations.gff3.gz |
| *Caenorhabditis bovis* | Cbov | 2654633 | https://ftp.ebi.ac.uk/pub/databases/wormbase/parasite/releases/WBPS16/species/caenorhabditis_bovis/PRJEB34497/caenorhabditis_bovis.PRJEB34497.WBPS16.genomic.fa.gz | https://ftp.ebi.ac.uk/pub/databases/wormbase/parasite/releases/WBPS16/species/caenorhabditis_bovis/PRJEB34497/caenorhabditis_bovis.PRJEB34497.WBPS16.annotations.gff3.gz |
| *Caenorhabditis brenneri* | Cbre | 135651 | https://ftp.ebi.ac.uk/pub/databases/wormbase/parasite/releases/WBPS16/species/caenorhabditis_brenneri/PRJNA20035/caenorhabditis_brenneri.PRJNA20035.WBPS16.genomic.fa.gz | https://ftp.ebi.ac.uk/pub/databases/wormbase/parasite/releases/WBPS16/species/caenorhabditis_brenneri/PRJNA20035/caenorhabditis_brenneri.PRJNA20035.WBPS16.annotations.gff3.gz |
| *Caenorhabditis briggsae* | Cbri | 6238 | https://ftp.ebi.ac.uk/pub/databases/wormbase/parasite/releases/WBPS16/species/caenorhabditis_briggsae/PRJNA10731/caenorhabditis_briggsae.PRJNA10731.WBPS16.genomic.fa.gz | https://ftp.ebi.ac.uk/pub/databases/wormbase/parasite/releases/WBPS16/species/caenorhabditis_briggsae/PRJNA10731/caenorhabditis_briggsae.PRJNA10731.WBPS16.annotations.gff3.gz |
| *Caenorhabditis elegans* | Cele | 6239 | https://ftp.ebi.ac.uk/pub/databases/wormbase/parasite/releases/WBPS16/species/caenorhabditis_elegans/PRJNA13758/caenorhabditis_elegans.PRJNA13758.WBPS16.genomic.fa.gz | https://ftp.ebi.ac.uk/pub/databases/wormbase/parasite/releases/WBPS16/species/caenorhabditis_elegans/PRJNA13758/caenorhabditis_elegans.PRJNA13758.WBPS16.annotations.gff3.gz |
| *Caenorhabditis inopinata* | Cino | 1978547 | https://ftp.ebi.ac.uk/pub/databases/wormbase/parasite/releases/WBPS16/species/caenorhabditis_inopinata/PRJDB5687/caenorhabditis_inopinata.PRJDB5687.WBPS16.genomic.fa.gz | https://ftp.ebi.ac.uk/pub/databases/wormbase/parasite/releases/WBPS16/species/caenorhabditis_inopinata/PRJDB5687/caenorhabditis_inopinata.PRJDB5687.WBPS16.annotations.gff3.gz |
| *Caenorhabditis japonica* | Cjap | 281687 | https://ftp.ebi.ac.uk/pub/databases/wormbase/parasite/releases/WBPS16/species/caenorhabditis_japonica/PRJNA12591/caenorhabditis_japonica.PRJNA12591.WBPS16.genomic.fa.gz | https://ftp.ebi.ac.uk/pub/databases/wormbase/parasite/releases/WBPS16/species/caenorhabditis_japonica/PRJNA12591/caenorhabditis_japonica.PRJNA12591.WBPS16.annotations.gff3.gz |
| *Caenorhabditis latens* | Clat | 1503980 | https://ftp.ebi.ac.uk/pub/databases/wormbase/parasite/releases/WBPS16/species/caenorhabditis_latens/PRJNA248912/caenorhabditis_latens.PRJNA248912.WBPS16.genomic.fa.gz | https://ftp.ebi.ac.uk/pub/databases/wormbase/parasite/releases/WBPS16/species/caenorhabditis_latens/PRJNA248912/caenorhabditis_latens.PRJNA248912.WBPS16.annotations.gff3.gz |
| *Caenorhabditis nigoni* | Cnig | 1611254 | https://ftp.ebi.ac.uk/pub/databases/wormbase/parasite/releases/WBPS16/species/caenorhabditis_nigoni/PRJNA384657/caenorhabditis_nigoni.PRJNA384657.WBPS16.genomic.fa.gz | https://ftp.ebi.ac.uk/pub/databases/wormbase/parasite/releases/WBPS16/species/caenorhabditis_nigoni/PRJNA384657/caenorhabditis_nigoni.PRJNA384657.WBPS16.annotations.gff3.gz |
| *Caenorhabditis panamensis* | Cpan | 2301259 | https://ftp.ebi.ac.uk/pub/databases/wormbase/parasite/releases/WBPS16/species/caenorhabditis_panamensis/PRJEB28259/caenorhabditis_panamensis.PRJEB28259.WBPS16.genomic.fa.gz | https://ftp.ebi.ac.uk/pub/databases/wormbase/parasite/releases/WBPS16/species/caenorhabditis_panamensis/PRJEB28259/caenorhabditis_panamensis.PRJEB28259.WBPS16.annotations.gff3.gz |
| *Caenorhabditis parvicauda* | Cpar | 2305859 | https://ftp.ebi.ac.uk/pub/databases/wormbase/parasite/releases/WBPS16/species/caenorhabditis_parvicauda/PRJEB12595/caenorhabditis_parvicauda.PRJEB12595.WBPS16.genomic.fa.gz | https://ftp.ebi.ac.uk/pub/databases/wormbase/parasite/releases/WBPS16/species/caenorhabditis_parvicauda/PRJEB12595/caenorhabditis_parvicauda.PRJEB12595.WBPS16.annotations.gff3.gz |
| *Caenorhabditis quiockensis* | Cqui | 2305861 | https://ftp.ebi.ac.uk/pub/databases/wormbase/parasite/releases/WBPS16/species/caenorhabditis_quiockensis/PRJEB11354/caenorhabditis_quiockensis.PRJEB11354.WBPS16.genomic.fa.gz | https://ftp.ebi.ac.uk/pub/databases/wormbase/parasite/releases/WBPS16/species/caenorhabditis_quiockensis/PRJEB11354/caenorhabditis_quiockensis.PRJEB11354.WBPS16.annotations.gff3.gz |
| *Caenorhabditis remanei* | Crem | 31234 | https://ftp.ebi.ac.uk/pub/databases/wormbase/parasite/releases/WBPS16/species/caenorhabditis_remanei/PRJNA577507/caenorhabditis_remanei.PRJNA577507.WBPS16.genomic.fa.gz | https://ftp.ebi.ac.uk/pub/databases/wormbase/parasite/releases/WBPS16/species/caenorhabditis_remanei/PRJNA577507/caenorhabditis_remanei.PRJNA577507.WBPS16.annotations.gff3.gz |
| *Caenorhabditis sinica* | Csin | 1550068 | https://ftp.ebi.ac.uk/pub/databases/wormbase/parasite/releases/WBPS16/species/caenorhabditis_sinica/PRJNA194557/caenorhabditis_sinica.PRJNA194557.WBPS16.genomic.fa.gz | https://ftp.ebi.ac.uk/pub/databases/wormbase/parasite/releases/WBPS16/species/caenorhabditis_sinica/PRJNA194557/caenorhabditis_sinica.PRJNA194557.WBPS16.annotations.gff3.gz |
| *Caenorhabditis sulstoni* | Csul | 2305862 | https://ftp.ebi.ac.uk/pub/databases/wormbase/parasite/releases/WBPS16/species/caenorhabditis_sulstoni/PRJEB12601/caenorhabditis_sulstoni.PRJEB12601.WBPS16.genomic.fa.gz | https://ftp.ebi.ac.uk/pub/databases/wormbase/parasite/releases/WBPS16/species/caenorhabditis_sulstoni/PRJEB12601/caenorhabditis_sulstoni.PRJEB12601.WBPS16.annotations.gff3.gz |
| *Caenorhabditis tribulationis* | Ctri | 2306311 | https://ftp.ebi.ac.uk/pub/databases/wormbase/parasite/releases/WBPS16/species/caenorhabditis_tribulationis/PRJEB12608/caenorhabditis_tribulationis.PRJEB12608.WBPS16.genomic.fa.gz | https://ftp.ebi.ac.uk/pub/databases/wormbase/parasite/releases/WBPS16/species/caenorhabditis_tribulationis/PRJEB12608/caenorhabditis_tribulationis.PRJEB12608.WBPS16.annotations.gff3.gz |
| *Caenorhabditis tropicalis* | Ctro | 1561998 | https://ftp.ebi.ac.uk/pub/databases/wormbase/parasite/releases/WBPS16/species/caenorhabditis_tropicalis/PRJNA53597/caenorhabditis_tropicalis.PRJNA53597.WBPS16.genomic.fa.gz | https://ftp.ebi.ac.uk/pub/databases/wormbase/parasite/releases/WBPS16/species/caenorhabditis_tropicalis/PRJNA53597/caenorhabditis_tropicalis.PRJNA53597.WBPS16.annotations.gff3.gz |
| *Caenorhabditis uteleia* | Cute | 2305860 | https://ftp.ebi.ac.uk/pub/databases/wormbase/parasite/releases/WBPS16/species/caenorhabditis_uteleia/PRJEB12600/caenorhabditis_uteleia.PRJEB12600.WBPS16.genomic.fa.gz | https://ftp.ebi.ac.uk/pub/databases/wormbase/parasite/releases/WBPS16/species/caenorhabditis_uteleia/PRJEB12600/caenorhabditis_uteleia.PRJEB12600.WBPS16.annotations.gff3.gz |
| *Caenorhabditis waitukubuli* | Cwai | 2305864 | https://ftp.ebi.ac.uk/pub/databases/wormbase/parasite/releases/WBPS16/species/caenorhabditis_waitukubuli/PRJEB12602/caenorhabditis_waitukubuli.PRJEB12602.WBPS16.genomic.fa.gz | https://ftp.ebi.ac.uk/pub/databases/wormbase/parasite/releases/WBPS16/species/caenorhabditis_waitukubuli/PRJEB12602/caenorhabditis_waitukubuli.PRJEB12602.WBPS16.annotations.gff3.gz |
| *Caenorhabditis zanzibari* | Czan | 2306312 | https://ftp.ebi.ac.uk/pub/databases/wormbase/parasite/releases/WBPS16/species/caenorhabditis_zanzibari/PRJEB12596/caenorhabditis_zanzibari.PRJEB12596.WBPS16.genomic.fa.gz | https://ftp.ebi.ac.uk/pub/databases/wormbase/parasite/releases/WBPS16/species/caenorhabditis_zanzibari/PRJEB12596/caenorhabditis_zanzibari.PRJEB12596.WBPS16.annotations.gff3.gz |
| *Caenorhabditis afra* | Cafr | 1094335 | http://download.caenorhabditis.org/v2/genome_files/CAFRA.caenorhabditis_afra_JU1286_v2.scaffolds.fna | http://download.caenorhabditis.org/v2/genome_files/CAFRA.caenorhabditis_afra_JU1286_v2.annotations.gff3 |
| *Caenorhabditis castelli* | Ccas | 1630362 | http://download.caenorhabditis.org/v2/genome_files/CCAST.caenorhabditis_castelli_JU1956_v2.scaffolds.fna | http://download.caenorhabditis.org/v2/genome_files/CCAST.caenorhabditis_castelli_JU1956_v2.annotations.gff3 |
| *Caenorhabditis doughertyi* | Cdou | 1094321 | http://download.caenorhabditis.org/v2/genome_files/CDOUG.caenorhabditis_doughertyi_JU1771_v2.scaffolds.fna | http://download.caenorhabditis.org/v2/genome_files/CDOUG.caenorhabditis_doughertyi_JU1771_v2.annotations.gff3 |
| *Caenorhabditis kamaaina* | Ckam | 1094325 | http://download.caenorhabditis.org/v2/genome_files/CKAMA.caenorhabditis_kamaaina_QG2077_v2.scaffolds.fna | http://download.caenorhabditis.org/v2/genome_files/CKAMA.caenorhabditis_kamaaina_QG2077_v2.annotations.gff3 |
| *Caenorhabditis macrosperma* | Cmac | 1094328 | http://download.caenorhabditis.org/v2/genome_files/CMACR.caenorhabditis_macrosperma_JU2083_v2.scaffolds.fna | http://download.caenorhabditis.org/v2/genome_files/CMACR.caenorhabditis_macrosperma_JU2083_v2.annotations.gff3 |
| *Caenorhabditis monodelphis* | Cmon | 1094320 | http://download.caenorhabditis.org/v2/genome_files/CMONO.caenorhabditis_monodelphis_JU1667_v2.scaffolds.fna | http://download.caenorhabditis.org/v2/genome_files/CMONO.caenorhabditis_monodelphis_JU1667_v2.annotations.gff3 |
| *Caenorhabditis nouraguensis* | Cnou | 1094327 | http://download.caenorhabditis.org/v2/genome_files/CNOUR.caenorhabditis_nouraguensis_JU2079_v2.scaffolds.fna | http://download.caenorhabditis.org/v2/genome_files/CNOUR.caenorhabditis_nouraguensis_JU2079_v2.annotations.gff3 |
| *Caenorhabditis plicata* | Cpli | 281681 | http://download.caenorhabditis.org/v2/genome_files/CPLIC.caenorhabditis_plicata_SB355_v2.scaffolds.fna | http://download.caenorhabditis.org/v2/genome_files/CPLIC.caenorhabditis_plicata_SB355_v2.annotations.gff3 |
| *Caenorhabditis virilis* | Cvir | 1094323 | http://download.caenorhabditis.org/v2/genome_files/CVIRI.caenorhabditis_virilis_JU1968_v2.scaffolds.fna | http://download.caenorhabditis.org/v2/genome_files/CVIRI.caenorhabditis_virilis_JU1968_v2.annotations.gff3 |
| *Caenorhabditis wallacei* | Cwal | 1094326 | http://download.caenorhabditis.org/v2/genome_files/CWALL.caenorhabditis_wallacei_JU1898_v2.scaffolds.fna | http://download.caenorhabditis.org/v2/genome_files/CWALL.caenorhabditis_wallacei_JU1898_v2.annotations.gff3 |

**Supplemental Table S1. Organisms' meta data.** For each of the seven focal model organisms, the list of the organisms used, including the scientific name, the abbreviated name used in the figures, the taxonomic ID, and the urls to the genomic FASTA and GFF3 annotation files.

| organism | rho | p-value |
|---|---|---|
| *Saccharomyces cerevisiae* | 0,77 | 7,20E-02 |
| *Homo sapiens* | 0,99 | 3,10E-04 |
| *Mus musculus* | 0,65 | 8,06E-02 |
| *Drosophila melanogaster* | 0,94 | 3,03E-11 |
| *Oryza sativa* | 0,90 | 1,60E-04 |
| *Arabidopsis thaliana* | 0,90 | 3,74E-02 |
| *Caenorhabditis elegans* | 0,80 | 5,71E-08 |

**Supplemental Table S2. Correlation between the conservation of the *de novo* emerged genes and the conservation of noncoding ORFs across the neighbor species.** For each focal species, the rho coefficient of the Spearman's rank correlation, along with the associated p-value.

| line name |
|---|
| Akaa, Finland (FI; lat. 61.10, long. 23.52, alt. 110, July 2018) |
| Lund, Sweden (SE; lat. 55.69, long. 13.19, alt. 28, August 2015) |
| Karensminde, Denmark (DK; lat. 55.94, long. 10.21, alt. 16, September 2014) |
| Gimenells, Spain (ES; lat. 41.66, long. 0.39, alt. 258, August 2018) |
| Siavonga, Zambia (ZI, Line ZI418; lat. -16.32, long. 28.42, alt. 479, July 2010) |
| Uman, Ukraine (UA; lat. 48.75, long. 30.21, alt. 214, August 2018) |
| Yesiloz, Turkey (TR; lat. 40.23, long. 32.26, alt. 680, September 2018) |

**Supplemental Table S3. *D. melanogaster* lines.** List of *D. melanogaster* lines used in Figure 5.
